# Functional Organization of the Maternal and Paternal Human 4D Nucleome

**DOI:** 10.1101/2020.03.15.992164

**Authors:** Stephen Lindsly, Wenlong Jia, Haiming Chen, Sijia Liu, Scott Ronquist, Can Chen, Xingzhao Wen, Cooper Stansbury, Gabrielle A. Dotson, Charles Ryan, Alnawaz Rehemtulla, Gilbert S. Omenn, Max Wicha, Shuai Cheng Li, Lindsey Muir, Indika Rajapakse

## Abstract

Every human somatic cell inherits a maternal and a paternal genome, which work together to give rise to cellular phenotypes. However, the allele-specific relationship between gene expression and genome structure through the cell cycle is largely unknown. By integrating haplotype-resolved genome-wide chromosome conformation capture, mature and nascent mRNA, and protein binding data, we investigate this relationship both globally and locally. We introduce the maternal and paternal 4D Nucleome, enabling detailed analysis of the mechanisms and dynamics of genome structure and gene function for diploid organisms. Our analyses find significant coordination between allelic expression biases and local genome conformation, and notably absent expression bias in universally essential cell cycle and glycolysis genes. We propose a model in which coordinated biallelic expression reflects prioritized preservation of essential gene sets.

## Introduction

Biallelic gene expression in diploid genomes inherently protects against potentially harmful mutations. Disrupted biallelic expression of certain genes increases vulnerability to disease in humans, such as in familial cancer syndromes that have loss of function in one allele (*1*). BRCA1 and BRCA2 are quintessential examples, for which missense, nonsense, or frameshift mutations affecting function of one allele significantly increase the risk of breast cancer in women (*2, 3*). Imprinted genes are also associated with multiple disease phenotypes such as Angelman and Prader-Willi syndromes (*4, 5)*. Other genes with monoallelic or allele-biased expression (MAE, ABE) may be associated with disease, but the contribution of allelic bias to disease phenotypes remains poorly understood.

ABE can occur with single nucleotide variants (SNVs), insertions or deletions (InDels), and chromatin modifications (*6–10*). Analyses of allelic bias suggest high variance across tissues and individuals, with estimates ranging from 4% to 26% of genes in a given setting (*7, 11*). In addition, higher order chromatin conformation and spatial positioning in the nucleus shape gene expression (*12–15*). As the maternal and paternal alleles can be distant in nucleus, their spatial positions may promote ABE (*16, 17*).

A major step towards understanding the contribution of allelic bias to disease is to identify ABE genes, recognizing that important biases may be transient and challenging to detect. Allele-specific expression and 3D structures are not inherently accounted for in genomics methods such as RNA-sequencing and genome-wide chromosome conformation capture (Hi-C). These limitations complicate interpretations of structure-function relationships, and complete phasing of the two genomes remains a significant challenge.

To improve understanding of ABE in genomic structure-function relationships, we developed a novel phasing algorithm for Hi-C data, which we integrate with allele-specific RNA-seq and Bru-seq data across three phases of the cell cycle in human B-lymphocytes (**Figure 1)**. RNA-seq and Bru-seq data were separated into their maternal and paternal components through SNVs/InDels (*18*). Our algorithm, HaploHiC, uses phased SNVs/InDels to impute Hi-C reads of unknown parental origin. Publicly available allele-specific protein binding data (ChIP-seq) were also included to better understand potential regulatory elements involved in allelic bias (*7, 19*). In addition to identifying known ABE genes silenced by X-Chromosome inactivation (XCI) or imprinting, our analyses find novel expression biases between alleles and cell cycle phases in several hundred genes, many of which had corresponding bias in allelespecific protein binding. Furthermore, the alleles of ABE genes were significantly more likely to differ in local structure compared to randomly selected alleles. In contrast, we observed a pronounced lack of ABE in crucial biological pathways and essential genes. Our findings highlight advantages of integrating genomics analyses in a cell cycle and allele-specific manner and represent an allele-specific extension of the 4D Nucleome (4DN) (*20–23*). This approach will be beneficial to investigation of human phenotypic traits and their penetrance, genetic diseases, vulnerability to complex disorders, and tumorigenesis.

**Fig. 1:**
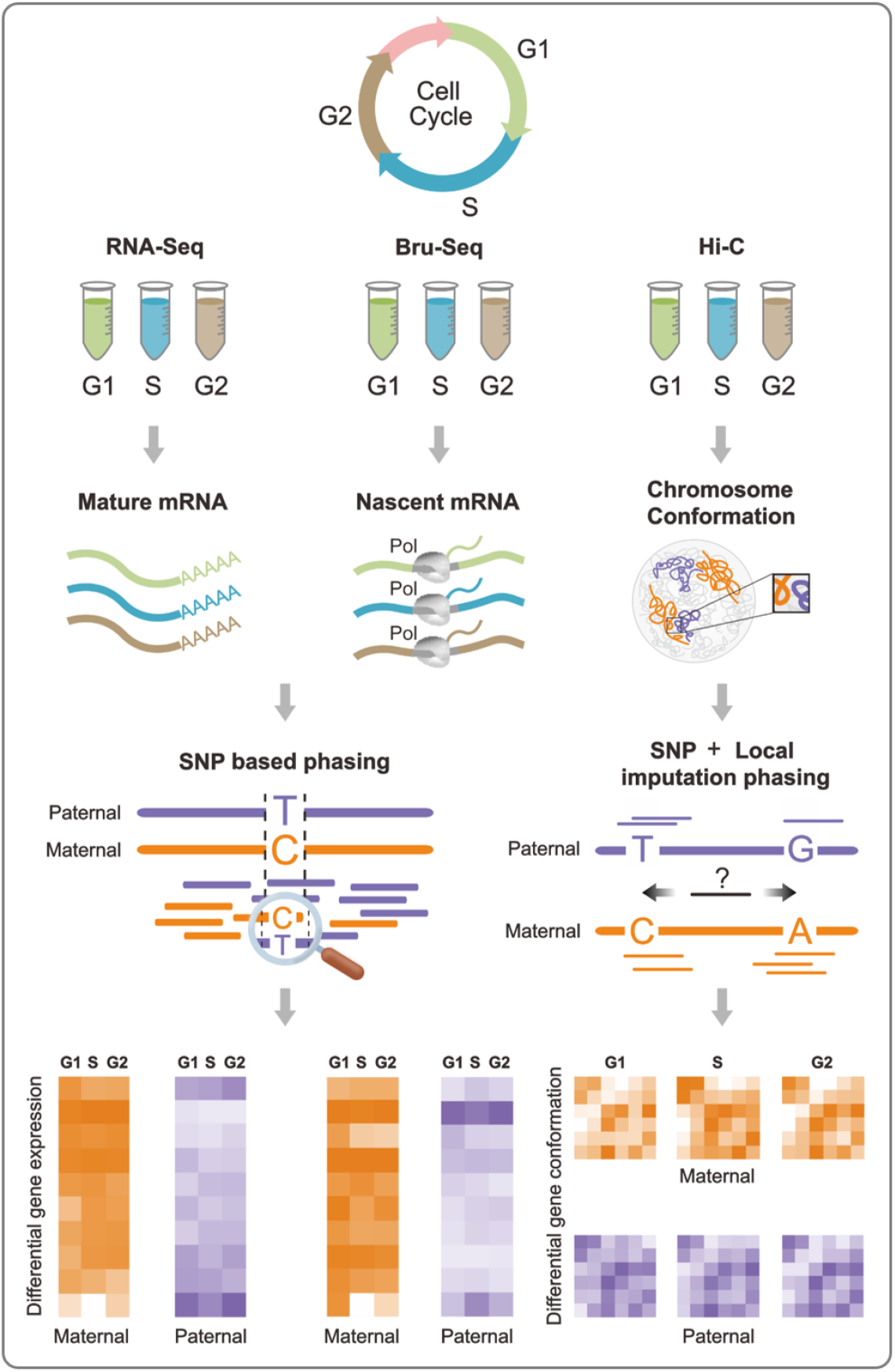
Experimental and allelic separation workflow. Cell cycle sorted cells were extracted for RNA-seq, Bru-seq, and Hi-C (left to right, respectively). RNA-seq and Bru-seq data were allelicly phased via SNVs/InDels (left). SNV/InDel based imputation and haplotype phasing of Hi-C data using HaploHiC (right). These data provide quantitative measures of structure and function of the maternal and paternal genomes through the cell cycle.

## Materials and Methods

### Cell Culture and Cell Cycle Sorting

Human GM12878 cells were cultivated in RPMI1640 medium supplemented with 10% fetal bovine serum (FBS). Live cells were stained with Hoechst 33342 (Cat #B2261, Sigma-Aldrich), and then sorted by fluorescence-activated cell sorting (FACS) to obtain cell fractions at the corresponding cell cycle phases G1, S, and G2/M (**Figure S2**).

### RNA-seq and Bru-seq Sequencing

Total RNA was extracted from sorted live cells for both RNA-seq and Bru-seq. We performed 5’-bromouridine (Bru) incorporation in live cells for 30 minutes, and the Bru-labeled cells were then stained on ice with Hoechst 33342 for 30 minutes before sorting at 4°C to isolate G1, S, and G2/M phase cells. The sorted cells were immediately lysed in TRizol (Cat # 15596026, ThermoFisher) and frozen. To isolate Bru-labeled RNA, DNAse-treated total RNA was incubated with anti-BrdU antibodies conjugated to magnetic beads (*24*). We converted the transcripts from the RNA-seq and Bru-seq experiments for all samples into cDNA libraries and deep-sequenced at 50-base length on an Illumina HiSeq2500 platform. The RNA-seq and Bru-seq data each consist of three biological replicates. From our RNA-seq replicates, we obtained a total of 193.4, 197.2, and 202.0 million raw reads for G1, S, and G2/M, respectively. From our Bru-seq replicates, we obtained a total of 162.5, 149.9, and 138.0 million raw reads for G1, S, and G2/M, respectively.

### Hi-C Sequencing

For cells used in construction of Hi-C libraries, cells were crosslinked with 1% formaldehyde, the reaction was neutralized with 0.125 M glycine, then cells were stained with Hoechst 33342 and sorted into G1, S, and G2/M fractions. Cross-linked chromatin was digested with the restriction enzyme MboI for 12 hours. The restriction enzyme fragment ends were tagged with biotin-dATP and ligated in situ. After ligation, the chromatins were de-cross-linked, and DNA was isolated for fragmentation. DNA fragments tagged by biotin-dATP, in the size range of 300-500 bp, were pulled down for sequencing adaptor ligation and polymerase chain reaction (PCR) products. The PCR products were sequenced on an Illumina HiSeq2500 platform. Respective to G1, S, and G2/M, we obtained 512.7, 550.3, and 615.2 million raw Hi-C sequence reads.

### RNA-seq and Bru-seq Data Processing

RNA-seq and Bru-seq analysis were performed as previously described (*25, 26*). Briefly, Bru-seq used Tophat (v1.3.2) to align reads without de novo splice junction calling after checking quality with FastQC (version 0.10.1). A custom gene annotation file was used in which introns are included but preference to overlapping genes is given on the basis of exon locations and stranding where possible (see (*26*) for full details). Similarly for RNA-seq data processing, the raw reads were checked with FastQC. Tophat (version 2.0.11) and Bowtie (version 2.1.0.0) were used to align the reads to the reference transcriptome (HG19). Cufflinks (version 2.2.1) was used for expression quantification, using UCSC hg19.fa and hg19.gtf as the reference genome and transcriptome, respectively. A locally developed R script using CummeRbund was used to format the Cufflinks output.

### Separation of Maternal and Paternal RNA-seq and Bru-seq Data

To determine allele-specific transcription and gene expression through Bru-seq and RNA-seq, all reads were aligned using GSNAP, a SNV aware aligner (*27, 28*). HG19 and UCSC gene annotations were used for the reference genome and gene annotation, respectively. The gene annotations were used to create the files for mapping to splice sites (used with −s option). Optional inputs to perform SNV aware alignment were also included. Specifically, −v was used to include the list of heterozygous SNVs and −use-sarray=0 was used to prevent bias against non-reference alleles (*29*).

After alignment, the output SAM files were converted to BAM files, sorted and indexed using SAMtools (*30*). SNV alleles were quantified using bam-readcounter to count the number of each base that was observed at each of the heterozygous SNV locations. Allele-specificity of each gene was then assessed by combining all of the SNVs in each gene. For RNA-seq, only exonic SNVs were used. Bru-seq detects nascent transcripts containing both exons and introns, so both exonic and intronic SNVs were used. Maternal and paternal gene expression were calculated by multiplying the genes’ overall read counts by the fraction of the SNV-covering reads that were maternal and paternal, respectively. We identified 266,899 SNVs from the Bru-seq data, compared with only 65,676 SNVs from RNA-seq data. However in the Bru-seq data, many SNVs have low read coverage depth. We required at least 5 SNV-covering reads for a SNV to be used to separate the maternal and paternal contributions to gene expression. This criterion found that there were similar numbers of informative SNVs (19,394 and 19,998) in the RNA-seq and Bru-seq data, respectively. Genes which did not contain informative SNVs were divided equally into their maternal and paternal contributions.

### Allele-specific Differential Expression

For a gene’s expression to be considered for differential expression analysis, we require each of the three replicates to have an average of at least 10 SNV-covering reads mapped to at least one of the alleles in all three cell cycle phases. This threshold was introduced to reduce the influence of technical noise on our differential expression results. From the 23,277 Refseq genes interrogated, there were 4,193 genes with at least 10 read counts mapped to either the maternal or paternal allele (or both) in the RNA-seq data. From Bru-seq, there were 5,294 genes using the same criterion. We refer to these genes as “allele-specific” genes for their respective data sources. We observed that there were larger variances between samples and lower read counts in the Bru-seq data set than in RNA-seq. We identified differentially expressed genes between alleles and between cell cycle phases for both RNA-seq and Bru-seq using a MATLAB implementation of DESeq (*31*). To reduce the possibility of false positives when determining differential expression, we imposed a minimum FPKM level of 0.1, a false discovery rate adjusted *p*-value threshold of 0.05, and a fold change cutoff of FC >2 for both RNA-seq and Bru-seq (*32*).

### Separation of Maternal and Paternal Hi-C Data by HaploHiC

Hi-C library construction and Illumina sequencing were performed using established methods (*9*). In this study, we separate the maternal and paternal genomes’ contributions to the Hi-C contact matrices to analyze their similarities and differences in genome structure. In order to determine which Hi-C reads come from which parental origin, we utilize differences in genomic sequence at phased SNVs/InDels. As these variations are unique to the maternal and paternal genomes, they can be used to distinguish reads. When attempting to separate the maternal and paternal genomes, complications arise when there are sections of DNA that are identical. There are a relatively small number of allele-specific variations, and the resulting segregated maternal and paternal contact matrices are sparse. In order to combat this problem, we seek to infer contacts of unknown parental origin.

We propose a novel technique, HaploHiC, for phasing reads of unknown parental origin using local imputation from known reads. HaploHiC uses a data-derived ratio based on the following hypothesis: if the maternal and paternal genomes have different 3D structures, we can use the reads with known origin (at SNV/InDel loci) to predict the origin of neighboring unknown reads (**Figure 4A, Supplemental Methods A**) (*33*). For example, if we observe that many contacts between two loci can be directly mapped to the paternal genome but few to the maternal genome, then unphased contacts between those loci are more likely to be from the paternal genome as well, and vice versa. This process of imputing Hi-C reads of unknown origin based on nearby known reads is similar to the methods developed by Tan *et al*. (*10*).

**Fig. 2:**
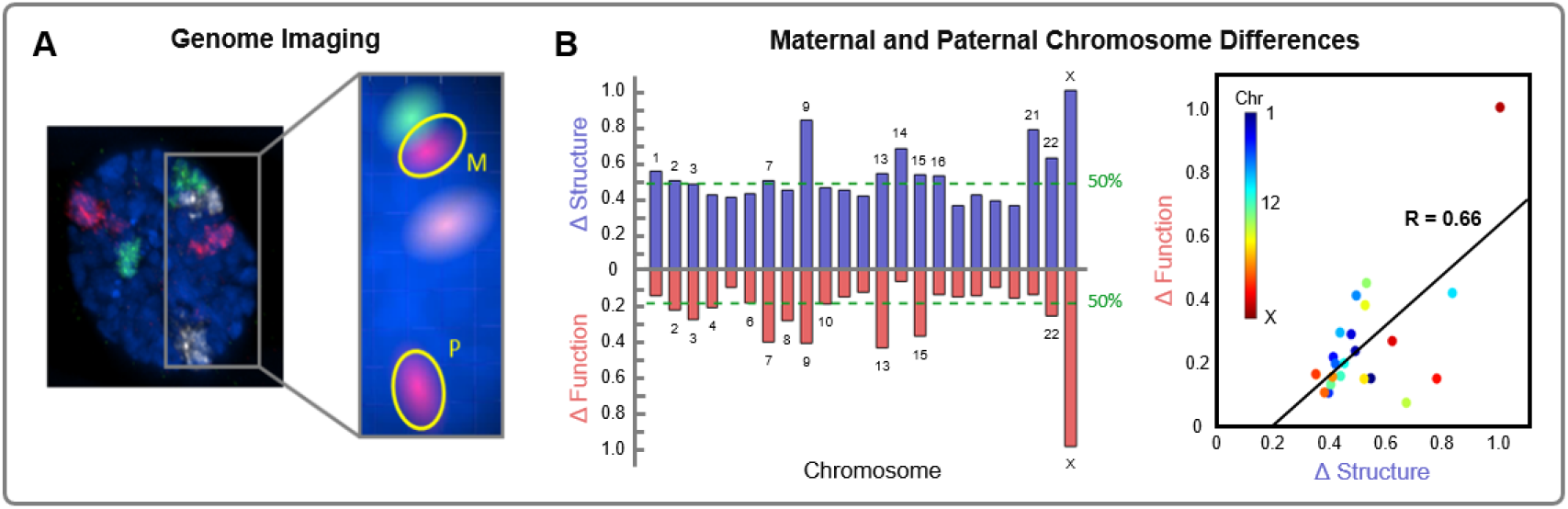
Genome imaging and chromosome differences. **(A)** Nucleus of a primary human fibroblast imaged using 3D FISH with the maternal and paternal copies of Chromosome 6, 8, and 11 painted red, green, and white, respectively (left). Subsection highlighting the separation between the maternal and paternal copies of Chromosome 11, now colored red (right). **(B)** Normalized chromosome level structural and functional parental differences of GM12878 cells. Structural differences (ΔStructure, blue) represent the aggregate changes between maternal and paternal Hi-C over all 1 Mb loci for each chromosome, adjusted for chromosome size in G1. Functional differences (ΔFunction, red) represent the aggregate changes between maternal and paternal RNA-seq over all 1 Mb loci for each chromosome, adjusted for chromosome size in G1. Green dashed lines correspond to the median structural (0.48, chromosome 3) and functional (0.20, chromosome 6) differences, in the top and bottom respectively, and all chromosomes equal to or greater than the threshold are labeled. Scatter plot of maternal and paternal differences in structure and function with best-fit line (R =0.66 and *p* < 0.05).

**Fig. 3:**
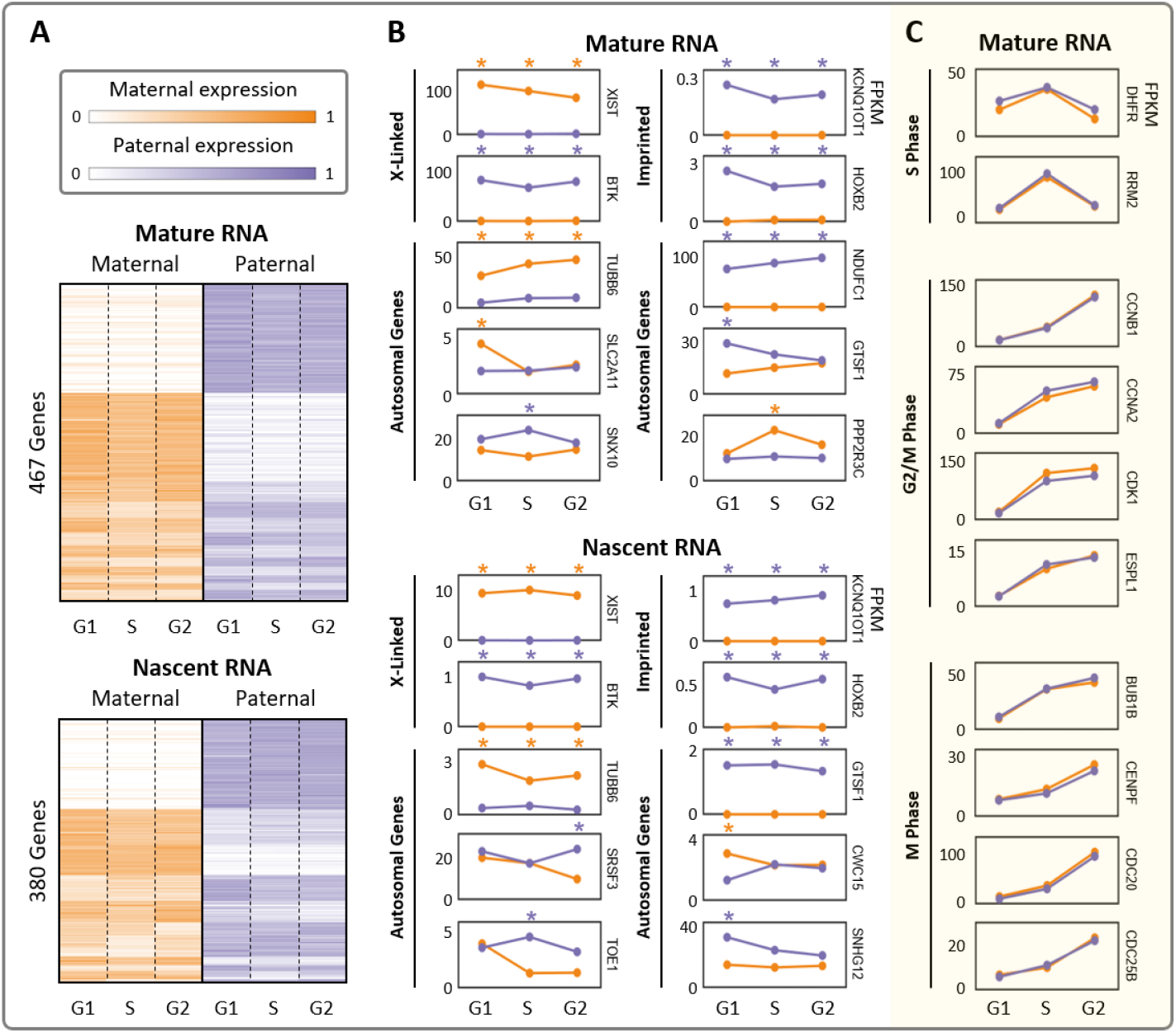
Allele-specific mature and nascent RNA expression. **(A)** Differentially expressed genes’ maternal and paternal RNA expression through the cell cycle. Expression heatmaps are average FPKM values over three replicates after row normalization. Genes are grouped by their differential expression patterns (**Figure S1**). **(B)** Representative examples of X-linked, imprinted, and other autosomal genes with allelic bias. Top and bottom sections of (A) and (B) show mature RNA levels (RNA-seq) and nascent RNA expression (Bru-seq), respectively. **(C)** Examples of cell cycle regulatory genes’ mature RNA levels through the cell cycle. These genes are grouped by their function in relation to the cell cycle and all exhibit CBE but none have ABE. All example genes in (B) and (C) reflect average FPKM values over three replicates, and ABE in a particular cell cycle phase is marked with an orange or purple asterisk for maternal or paternal bias, respectively. G2 includes both G2 and M phase.

**Fig. 4:**
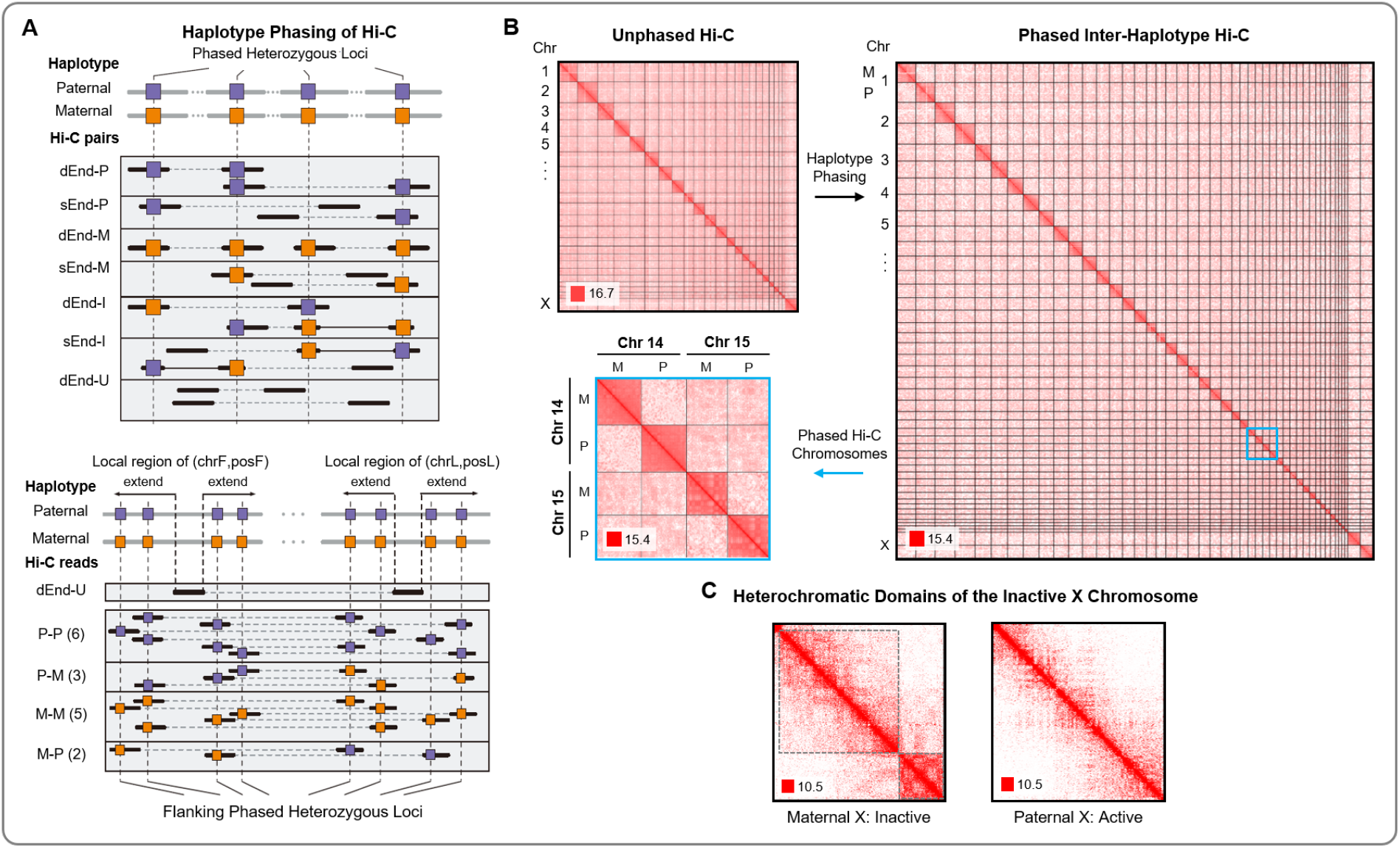
Haplotype phasing of Hi-C data. **(A)** HaploHiC separates paired-end reads into groups based on parental origin determined through SNVs/InDels (left, **Supplemental Methods A.3**). Reads are grouped by: (i) reads with one (sEnd-P/M) or both ends (dEnd-P/M) mapped to a single parent, (ii) reads are inter-haplotype, with ends mapped to both parents (d/sEnd-I), and (iii) reads with neither end mapped to a specific parent (dEnd-U). An example of a paired-end read (dEnd-U) with no SNVs/InDels has its origin imputed using nearby reads (right, **Supplemental Methods A.6**). A ratio of paternally and maternally mapped reads is found in a dynamically sized flanking region around the haplotype-unknown read’s location (**Supplemental Methods A.4**). This ratio then determines the likelihood of the haplotype-unknown read’s origin. **(B)** Whole-genome Hi-C of GM12878 cells (top left). Inter- and intra-haplotype chromatin contacts after phasing Hi-C data using HaploHiC (right). Chromosomes 14 and 15 highlight inter- and intra-chromosome contacts within and between genomes (bottom left). Visualized in *log_2_* scale 1 Mb resolution in G1. **(C)** Haplotype phasing illustrates that the inactive maternal Chromosome X is partitioned into large heterochromatic domains, outlined in dotted black boxes. Visualized in *log_2_* scale 100 kb resolution in G1.

HaploHiC marks paired-end reads as haplotype-known or -unknown depending on their coverage of heterozygous phased SNVs/InDels. Haplotype-known reads are directly assigned to their corresponding haplotype, maternal or paternal. HaploHiC uses a local contacts-based algorithm to impute the haplotype of haplotype-unknown reads using nearby SNVs/InDels. If the minimum threshold (ten paired-ends) of haplotype-known reads for local imputation is not reached, HaploHiC randomly assigns the haplotype-unknown reads to be maternal or paternal (less than 5% of all haplotype-unknown reads). Detailed materials and methods for haplotype phasing and Hi-C construction are provided in **Supplemental Methods A**.

Our validation shows that HaploHiC performs well, with an average accuracy of 96.9%, 97.2%, and 97.3% for G1, S, and G2, respectively, over 10 trials each (**Table S12**). Each validation trial randomly removed 10% of the heterozygous phased SNVs/InDels, and calculated imputation accuracy by the fraction of correctly imputed reads from the haplotype-known Hi-C reads covering these removed heterozygous mutations (Supplemental Methods A.8). Our validation of imputation accuracy is similar to the method presented in Tan *et al*. (*10*). We also perform multiple simulations for further validation (Supplemental Methods A.8). HaploHiC is available through a GitHub repository.

After haplotype assignment through HaploHiC, Hi-C paired-end reads (PE-reads) were distributed to intra-haplotype (P-P and M-M) and inter-haplotype (P-M and M-P) (**Supplemental Methods A.1-A.8**). Juicer was applied on intra-haplotype PE-reads, and outputs maternal and paternal contact matrices which were normalized through the Knight-Ruiz method of matrix balancing (*34, 35*). Inter-haplotype contact matrices were generated by HaploHiC (**Supplemental Methods A.7**). Intra- and inter-haplotype contacts are shown in **Figure 4B** and **Figure S8**. Both base pair level and fragment level matrices were constructed. The resolution of base pair level matrices are 1 Mb and 100 kb. Gene-level contacts were converted from fragment level matrices by HaploHiC.

## Results

### Chromosome-Scale Maternal and Paternal Differences

Spatial positioning of genes within the nucleus is known to be associated with transcriptional status (*12–15*). One might expect that the maternal and paternal copies of each chromosome would stay close together to ensure that their respective alleles have equal opportunities for transcription. Imaging of chromosome territories has shown that this is often not the case (**Figure 2A**, **Supplemental Methods B**) (*36–38*). This observation inspired us to investigate whether the two genomes operate in a symmetric fashion, or if allele-specific differences exist between the genomes regarding their respective chromatin organization patterns (structure) and gene expression profiles (function). We analyzed parentally phased whole-chromosome Hi-C and RNA-seq data at 1 Mb resolution to identify allele-specific differences in structure and function, respectively. We subtracted each chromosome’s paternal Hi-C matrix from the maternal matrix and found the Frobenius norm of the resulting difference matrix. The Frobenius norm provides a measure of distance between matrices, where equivalent maternal and paternal genome structures would result in a value of zero. Similarly, we subtracted the phased RNA-seq vectors in *log*_2_ scale and found the Frobenius norm of each difference vector. The Frobenius norms were adjusted for chromosome size and normalized for both Hi-C and RNA-seq.

We found that all chromosomes have allelic differences in both structure (Hi-C, blue) and function (RNA-seq, red) (**Figure 2B**). Chromosome X had the largest structural difference, as expected, followed by Chromosomes 9, 21, and 14. Chromosome X had the most extreme functional differences as well, followed by Chromosomes 13, 7, and 9. A threshold was assigned at the median Frobenius norm for Hi-C and RNA-seq (**Figure 2B** green dashed lines). The majority of chromosomes with larger structural differences than the median in Hi-C also have larger functional differences than the median in RNA-seq. There is a positive correlation between chromosome level differences in structure and function, which is statistically significant only when Chromosome X is included (R =0.66 and *p* < 0.05 with Chromosome X; R =0.30 and *p* =0.17 without Chromosome X).

### Allele-Specific RNA Expression

After confirming allelic differences in RNA expression at the chromosomal scale, we examined allele-specific expression of individual genes through RNA-seq and Bru-seq. We hypothesized that the chromosome scale expression differences were not only caused by known cases of ABE such as XCI and imprinting, but also by widespread ABE over many genes (*39, 40*). Therefore, we evaluated all allele-specific genes (genes with sufficient reads covering SNVs/InDels) for differential expression across the six settings: maternal and paternal in G1, S, and G2/M (hereafter, G2). These settings give rise to seven comparisons which consist of maternal versus paternal within each of the cell cycle phases (three comparisons), as well as G1 versus S and S versus G2 for the maternal and paternal genomes, respectively (two comparisons for each genome).

First, we identified genes with ABE and cell cycle-biased expression (CBE) from RNA-seq. While ABE refers to differential expression between alleles in each cell cycle phase, CBE refers to significant changes in expression from one cell cycle phase to another in each allele. From 23,277 RefSeq genes interrogated, there were 4,193 genes with sufficient coverage on SNVs/InDels to reliably determine allele-specific expression (*41*). We performed differential expression analysis for the seven comparisons to identify which of the 4,193 genes had ABE or CBE (*31*). We identified 615 differentially expressed genes from RNA-seq: 467 ABE genes, 229 CBE genes, and 81 genes with both ABE and CBE (**Tables S2**, **S4**). Both exons and introns containing informative SNVs/InDels were used for our Bru-seq data, from which 5,294 genes had sufficient coverage. We identified 505 differentially expressed genes from Bru-seq: 380 ABE genes, 164 CBE genes, and 39 genes with both ABE and CBE (**Tables S3**, **S5**). We also identified 130 genes that had ABE in both RNA-seq and Bru-seq. While this is substantially smaller than total number of ABE genes for RNA-seq and Bru-seq (467 and 380, respectively), the number of genes that are allele-specific in both data modalities is also smaller. That is, only 285 of the ABE genes from RNA-seq are allele-specific in Bru-seq and 192 of the ABE genes from Bru-seq are allele-specific in RNA-seq. The remaining genes did not have sufficient expression or SNV coverage to be included in the downstream analysis. We then separated the differentially expressed genes into their respective chromosomes to observe their distribution throughout the genome. From RNA-seq (Bru-seq), we found that autosomes had 3-14% ABE (1-11%) in their allele-specific genes which is comparable to previous findings (*11*). As expected, Chromosome X had a particularly high percentage of ABE genes at 90% (91%).

We identified 288 genes that have ABE in all three cell cycle phases from RNA-seq (160 paternally biased, 128 maternally biased) and 173 from Bru-seq (129 paternally biased, 44 maternally biased). This is the most common differential expression pattern among ABE genes and these genes form the largest clusters in **Figure 3A**. These clusters include, but are not limited to, XCI, imprinted, and other MAE genes. Known examples within these clusters are highlighted in the ‘X-Linked’ and ‘Imprinted’ sections of **Figure 3B**. We also identified hundreds of genes that are not currently appreciated in literature to have ABE, with examples shown in the ‘Autosomal Genes’ sections of **Figure 3B** for both mature and nascent RNA. Approximately half of all ABE genes were only differentially expressed in one or two cell cycle phases, which we refer to as transient allelic biases. These genes form the smaller clusters seen in **Figure 3A**. Examples of genes with transient allelic biases are also presented in the ‘Autosomal Genes’ section of **Figure 3B**. Transient expression biases like these may be due to coordinated expression of the two alleles in only certain cell cycle phases, though the mechanism behind this behavior is unclear.

Among the ABE genes from RNA-seq analysis, we found 117 MAE genes. In addition to the requirements for differential expression, we impose the thresholds of a FC ≥ 10 and for the inactive allele to have <0.1 Fragments Per Kilobase of transcript per Million (FPKM), or FC ≥ 50 across all three cell cycle phases. Our analysis confirmed MAE for imprinted and XCI genes, with examples shown in **Figure 3B**. Imprinted and XCI genes are silenced via transcriptional regulation, which was verified by monoallelic nascent RNA expression (Bru-seq). The *XIST* gene, which is responsible for XCI, was expressed in the maternal allele reflecting the deactivation of the maternal Chromosome X. XCI was also observed from Hi-C through large heterochromatic domains in the maternal Chromosome X, and the absence of these domains in the paternal Chromosome X (**Figure 4C**). The inactive Chromosome X in our cells is opposite of what is commonly seen for the GM12878 cell line in literature (likely due to our specific GM12878 sub-clone), but is consistent between our data modalities (*9, 10, 19*). The MAE genes also include six known imprinted genes, four expressed from the paternal allele (KCNQ1OT1, SNRPN, SNURF, and PEG10) and two from the maternal allele (NLRP2 and HOXB2). Some of the known imprinted genes that were confirmed in our data are associated with imprinting diseases, such as Beckwith-Wiedemann syndrome (KCNQ1OT1 and NLRP2), Angelman syndrome (SNRPN and SNURF), and Prader-Willi syndrome (SNRPN and SNURF) (*42, 43*). These genes and their related diseases offer further support for allele-specific analysis, as their monoallelic expression could not be detected otherwise.

After observing that approximately half of all ABE genes had transient expression biases, we hypothesized that alleles may have unique dynamics through the cell cycle. We then focused our investigation on allele-specific gene expression through the cell cycle to determine if alleles had CBE, and whether alleles were coordinated in their cell-cycle dependent expression (**Figure S1B**). We compared the expression of each allele between G1 and S as well as between S and G2, which provides insight into the differences between maternal and paternal alleles’ dynamics across the cell cycle. In the G1 to S comparison, there are 88 (*55*) genes in RNA-seq (Bru-seq) which have similar expression dynamics in both alleles. These genes’ maternal and paternal alleles are similarly upregulated or downregulated from G1 to S. In contrast, 87 (97) genes in RNA-seq (Bru-seq) have different expression dynamics between alleles. That is, only one allele is up or downregulated in the transition from G1 to S. In the S to G2 comparison, there are 26 (*3*) genes in RNA-seq (Bru-seq) that have similar expression dynamics in both alleles and 56 (*12*) genes with different expression dynamics between alleles. From these data, we see a coordination of expression between many, but certainly not all, alleles through the cell cycle.

### Biallelic Expression and Cellular Function

We observed from our analysis of CBE that multiple cell cycle regulatory genes had no instances of ABE (**Figure 3C**). We expanded this set of genes to include all allele-specific genes contained in the KEGG cell cycle pathway (*44*). Again, we found zero instances of ABE. This may suggest that genes with certain crucial cellular functions, like cell cycle regulation, may have coordinated biallelic expression to ensure their sufficient presence as a means of robustness. This is supported by previous findings which showed restricted genetic variation of enzymes in the essential glycolytic pathway (*45*).

We hypothesized that genes implicated in critical cell cycle processes would be less likely to have ABEs. We tested additional modules derived from KEGG pathways containing at least five allele-specific genes, with the circadian rhythm module supplemented by a known core circadian gene set (*20*). Examples of modules with varying proportions of ABE are shown in **Table 1** (**Table S7**), where “Percent ABE” refers to the proportion of genes with ABE to the total number of allele-specific genes in that module. We found that there are multiple crucial modules, including the glycolysis/gluconeogenesis and pentose phosphate pathways, which also had zero ABE genes. To explore the possibility of a global phenomenon by which genes essential to cellular fitness are significantly less likely to have biased expression, we analyzed the frequency of ABE in 1,734 genes experimentally determined to be essential in human cells (*46*). Using the 662 allele-specific genes in this set, we found that these essential genes were significantly less likely to have ABE than a random selection of allele-specific genes (5.8% versus 11.1%, p < 0.001, **Supplemental Methods D**), consistent with our hypothesis that critical genes are likely to be expressed by both alleles. In total, we offer a model in which coordinated biallelic expression reflects prioritized preservation of essential gene sets.

**Table 1:**
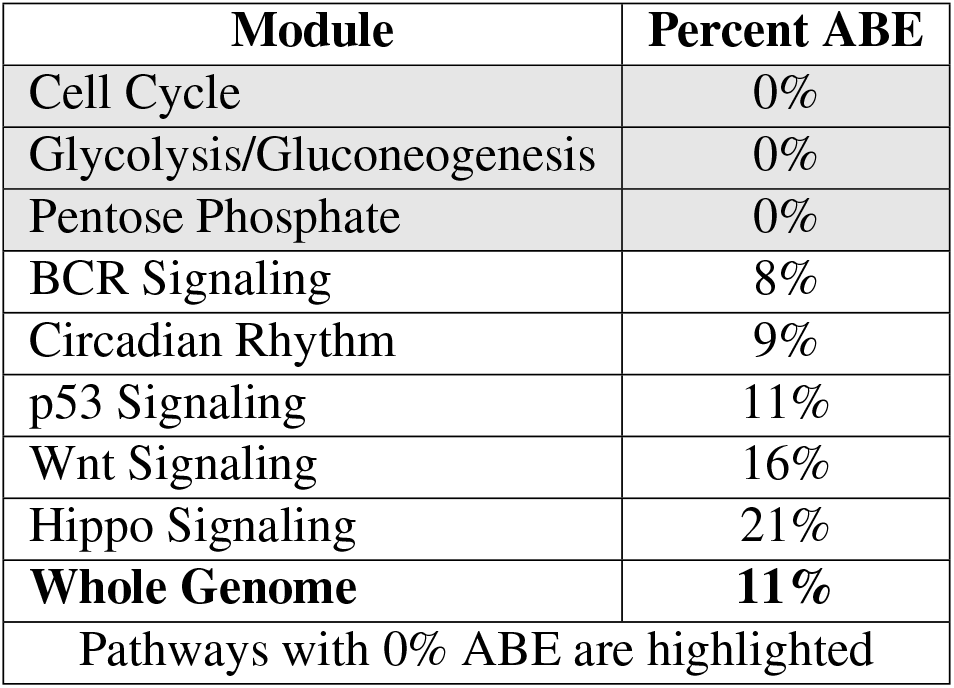
Allelic bias in biological modules.

| **Module** | **Percent ABE** |
| --- | --- |
| Cell Cycle | 0% |
| Glycolysis/Gluconeogenesis | 0% |
| Pentose Phosphate | 0% |
| BCR Signaling | 8% |
| Circadian Rhythm | 9% |
| p53 Signaling | 11% |
| Wnt Signaling | 16% |
| Hippo Signaling | 21% |
| **Whole Genome** | **11%** |
| Pathways with 0% ABE are highlighted |  |

### Allele-Specific Genome Structure

Motivated by our observations of chromosome level structural differences between the maternal and paternal genomes, we examined the HaploHiC separated data in more detail to determine where these differences reside. The genome is often categorized into two compartments: transcriptionally active euchromatin and repressed heterochromatin. In studies comparing multiple types of cells or cells undergoing differentiation, areas of euchromatin and heterochromatin often switch corresponding to genes that are activated/deactivated for the specific cell type (*47*). We explored this phenomenon in the context of the maternal and paternal Hi-C matrices to determine if the two genomes had differing chromatin compartments. Chromatin compartments can be identified from Hi-C data using methods such as principal component analysis or spectral clustering (**Supplemental Methods C**) (*48*). We applied spectral clustering to every chromosome across all three cell cycle phases at 1 Mb resolution. We found that there were slight changes in chromatin compartments for all chromosomes, but the vast majority of these changes took place on the borders between compartments rather than an entire region switching compartments. These border differences were not enriched for ABE genes. This implies that, although the structures may not be identical, the maternal and paternal genomes have similar overall compartmentalization (aside from Chromosome X).

We next applied spectral clustering recursively to the Hi-C data at 100 kb resolution to determine whether there were differences in TADs between the two genomes throughout the cell cycle (*48*). While the current understanding of genomic structure dictates that TAD boundaries are invariant (between alleles, cell types, etc), it is also known that “intra-TAD” structures are highly variable (*47, 49, 50*). The spectral identification method has an increased ability to discern these subtle structural changes. We found that TAD boundaries were variable between the maternal and paternal genomes and across cell cycle phases in all chromosomes. This supports previous findings of allelic differences in TADs for single cells, and we predict that they are even more variable across cell types (*48, 50*). Differences in TAD boundaries were observed surrounding MAE genes, ABE genes, and genes with coordinated biallelic expression (**Figure S6**). This indicated that changes in TAD boundaries were not directly related to allelic expression differences.

Although we did not find a direct relationship between TAD boundary differences and ABE genes, we observed during this analysis that the local genome structure around the six imprinted genes had noticeable differences. We then sought to analyze all genes with ABE or CBE to find out if they had corresponding structural differences at a local level. We analyzed the local Hi-C matrices for each of the 615 RNA-seq and 505 Bru-seq differentially expressed genes. Using a 300 kb flanking region centered at the 100 kb bin containing the transcription start site, we isolated a 7×7 matrix (700 kb) for each differentially expressed gene (**Figure 5A, B**). These matrices represent the local genomic structure of the differentially expressed genes, and are slightly smaller than average TAD size (~1 Mb). We then compared the correlation matrices of the *log*_2_-transformed local Hi-C data and determined whether or not the matrices have statistically significant differences (p < 0.05) (*23, 51*). We applied this comparison to all genes that were differentially expressed in RNA-seq (Bru-seq) and found that 515 (403) genes had at least one comparison in which both the expression and local structure had significantly changed. While changes in local genome structure and changes in gene expression do not have a one-to-one relationship, we found that both ABE and CBE genes are more likely to have significant architectural differences than randomly sampled allele-specific genes (p < 0.01) (**Figure S7** and **Supplemental Methods D**). This lends further support to the idea that there is a relationship between allele-specific differences in gene expression and genome structure.

**Fig. 5:**
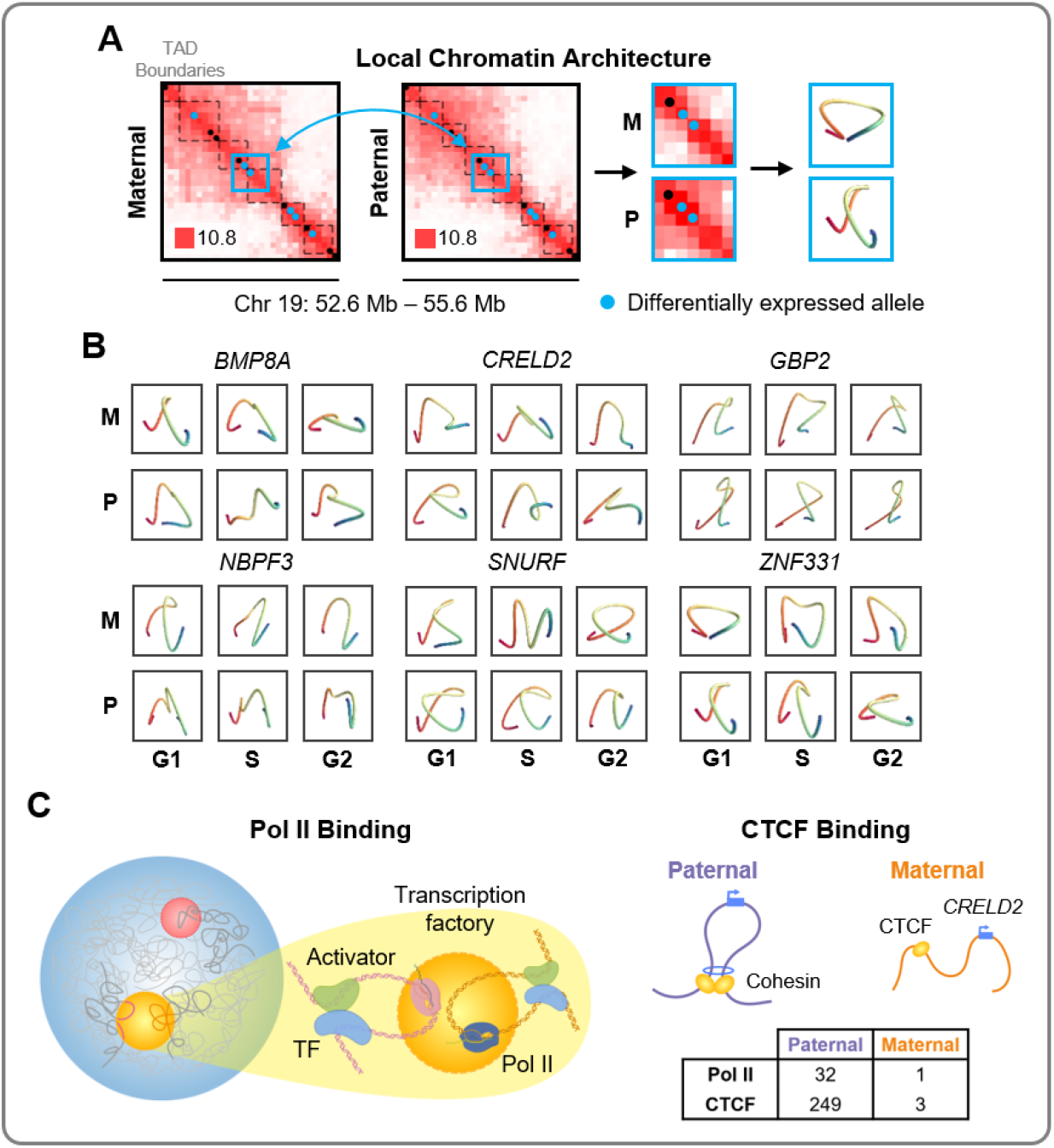
Local chromatin structure and transcription factor binding. **(A)** Local regions around differentially expressed genes are tested for significant conformational changes. These regions are modeled to visualize the conformations around each allele through G1, S, and G2 (**Supplemental Methods F**). Example of local chromatin structure extraction is shown for ZNF331 in G1 phase (center of blue box). Hi-C matrices are shown in *log_2_* scale 100 kb resolution. **(B)** 3D models of the local genome structure around six ABE genes with bias agreement of Pol II and significant changes in local genome structure. **(C)** Schematic representation of allele-specific Pol II and CTCF binding, with highlighted gene CRELD2, which had binding biases in both. Table shows extreme binding biases of Pol II and CTCF on CRELD2 as an example.

### Allele-Specific Protein Binding

To uncover the mechanisms behind the relationship between allele-specific gene expression and genome structure, we looked to DNA binding proteins such as RNA polymerase II (Pol II), CCCTC-binding factor (CTCF), and 35 other transcription factors. We used publicly available protein binding data from AlleleDB in tandem with RNA-seq and found 114 genes that have an allelic bias in both gene expression and binding of at least one such protein (*19*). We identified 13 genes which have ABE and biased binding of Pol II, with bias agreement in 11 cases (85%). That is, the allelic expression and Pol II binding were biased toward the same allele. For CTCF, 33 of 72 cases have bias agreement (46%), and for all other transcription factors analyzed, 20 of 29 cases have bias agreement (69%) (**Table S6**). The CTCF binding bias agreement of around 50% is expected, based on previous studies (*7*). This is likely due to CTCF’s role as an insulator, since an allele could be expressed or suppressed by CTCF’s presence depending on the context. To avoid potential inconsistencies between our data and the protein data from AlleleDB, we excluded Chromosome X when testing for ABE and protein binding biases.

We evaluated the relationship between TAD boundary differences between the maternal and paternal genomes and allele-specific CTCF binding sites. We found multiple instances of biased binding of CTCF and corresponding changes to the boundaries of TADs containing ABE genes. Examples of this phenomenon are shown in the center of Figure S6A, where TAD boundaries from the maternal (paternal) Hi-C data are closer to a maternally (paternally) biased CTCF binding site in some cell cycle phases near the ABE genes ANKRD19P, C9orf89, and FAM120A. Despite observing individual instances of biased CTCF binding corresponding to TAD boundary differences and ABE genes, there were insufficient data to evaluate this relationship genome-wide. We hypothesize that differences in TAD boundaries would correspond to allele-biased CTCF binding provided that there were enough data, as it has been repeatedly shown that TAD boundaries are enriched with CTCF binding (*49, 52)*.

We analyzed the 11 genes with allelic expression and Pol II binding bias agreement further to determine if they also had significant changes in local genome structure. Through local Hi-C comparisons, we found that all 11 of these genes had significant changes in structure in at least one cell cycle phase. 3D models for six of these genes are shown in **Figure 5B**, which highlight differences in local genome structure (**Supplemental Methods F**) (*53*). The genes with bias agreement and changes in local genome structure include known imprinted genes such as SNURF and SNRPN, as well as genes with known allele-specific expression (and suggested imprinting in other cell types) like ZNF331 (*54*). Additionally, there are multiple genes with known associations with diseases or disorders such as BMP8A, CRELD2, and NBPF3 (55–57). These findings suggest that changes in local structure often coincide with changes in expression due to the increased or decreased ability of a gene to access the necessary transcriptional machinery within transcription factories (*14, 58*). We visualize this relationship for the gene CRELD2 as an example (**Figure 5C**).

### The Maternal and Paternal 4D Nucleome

We define the maternal and paternal 4DN as the integration of allele-specific genome structure with gene expression data through time, adapted from Chen et al. (*20*). Many complex dynamical systems are investigated using a network perspective, which offers a simplified representation of a system (*59, 60)*. Networks capture patterns of interactions between their components and how those interactions change over time (*20*). We can consider genome structure as a network, since Hi-C data captures interactions between genomic loci (*12, 15*). In network science, the relative importance of a node in a network is commonly determined using network centrality (*59*). For Hi-C data, we consider genomic loci as nodes and use network centrality to measure the importance of each locus at each cell cycle phase (*23, 61*). We initially performed a global analysis of the maternal and paternal 4DN by combining RNA-seq with multiple network centrality measures (**Supplemental Methods D**) (*23*). We found differences between the maternal and paternal genomes and across cell cycle phases, but only Chromosome X had clear maternal and paternal separation (**Figures S9**, **S10**).

In our earlier analysis, we found a significant relationship between ABE and changes in local genome structure. We also observed that genes in multiple critical biological modules had coordinated biallelic expression. Motivated by these results, we performed an integrated analysis of structure and function to determine allele-specific dynamics of targeted gene sets. We constructed a sub-network for each gene set (analogous to an in silico 5C matrix), by extracting rows and columns of the Hi-C matrix containing genes of interest for each cell cycle phase (*62*). We used eigenvector centrality (similar to Google’s PageRank) to quantify structure, and used the average expression from the three RNA-seq replicates to quantify function, for each allele in the sub-network (*63*). We utilized the concept of a phase plane to plot the maternal and paternal 4DN (4DN phase plane, adapted from Chen et al.) (**Figure 6)** (*20*). We designated one axis as a measure of structure and the other as a measure of function. Coordinates of each point in the 4DN phase plane were determined from normalized structure data (x-axis, sub-network eigenvector centrality) and function data (*y*-axis, FPKM). The 4DN phase plane contains three points for each allele, which represent G1, S, and G2. We define allelic divergence (AD) as the average Euclidean distance between the maternal and paternal alleles across all cell cycle phases in the 4DN phase plane (**Supplemental Methods D**).

**Fig. 6:**
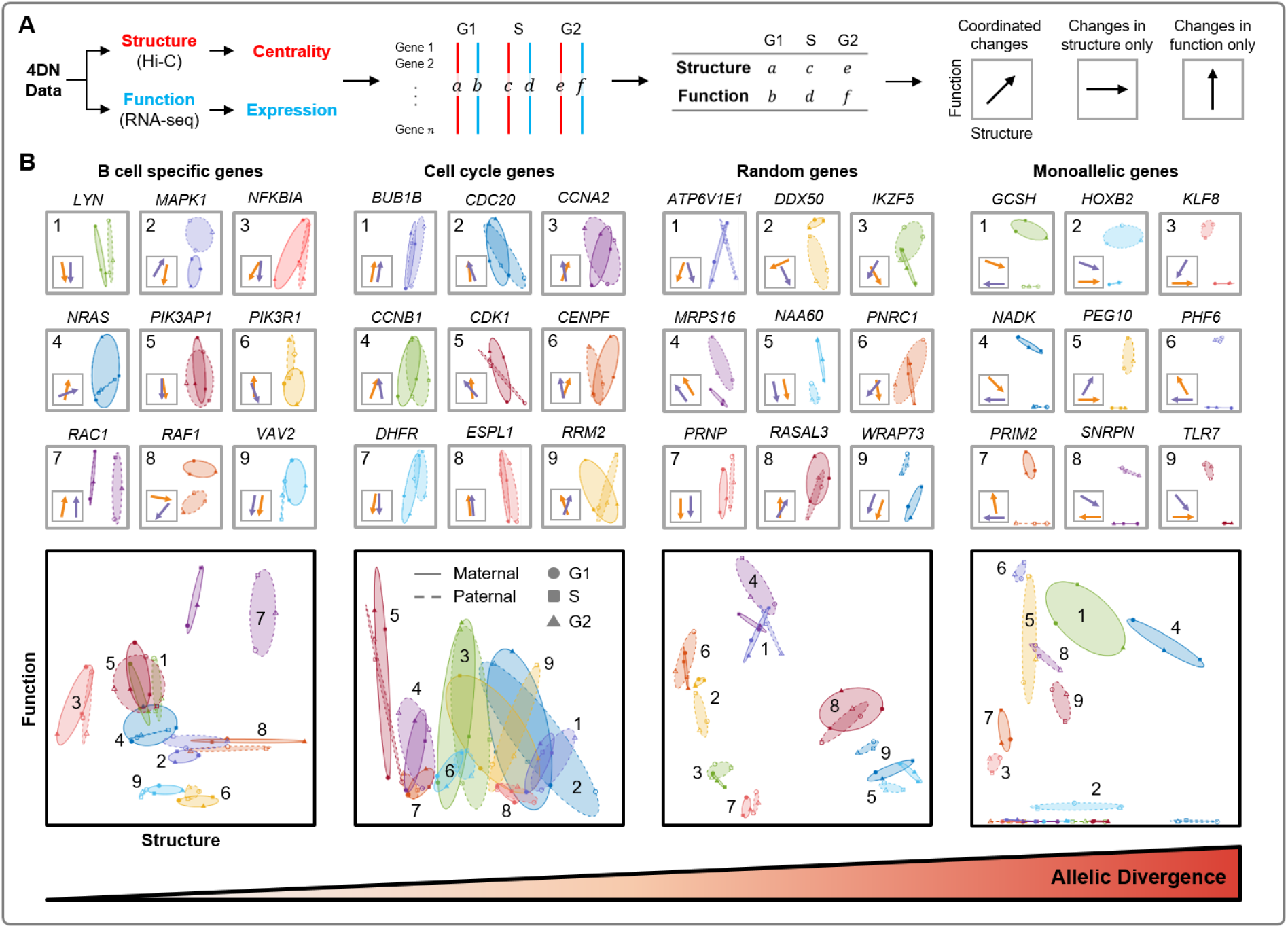
4DN phase planes reveal a wide range of allelic divergences in gene sub-networks. (**A**) Workflow to obtain structure and function measures. Eigenvector centrality for each gene is computed from the extracted sub-network of Hi-C contacts. Expression for each gene can be found directly from RNA-seq. Simplified phase planes are shown with linear relationships between changes in structure and function, changes in structure with no changes in function, and changes in function with no changes in structure. (**B**) 4DN phase planes of genes specific to B cell function, cell cycle genes, random allele-specific genes, and MAE genes, highlighting the similarities and differences between their alleles. Genes such as *BUB1B* and *PIK3AP1* have similar phase planes between alleles, while *RAC1* differs in structure and *WRAP73* differs in function. The bottom plot for each column combines the phase planes of the nine example genes, and the average allelic divergence is calculated from each of these gene sets.

We show four example 4DN phase planes of gene sub-networks with various ADs in **Figure 6.** Genes which are known to be crucial for cell cycle regulation have a mean AD of 0.0245 (**Figure 6B**, middle-left). Given that GM12878 is a B-lymphocyte cell line, we were interested in the AD of genes which are related to B cell receptor functionality. We found that these genes had a mean AD of 0.0225 (**Figure 6B**, left). The ADs of cell cycle regulating genes and B cell specific genes are smaller than the mean AD of randomly selected allele-specific genes (AD =0.0301 over 10,000 samples). This may be indicative of a robust coordination between the alleles to maintain proper cellular function and progression through the cell cycle, and therefore a lack of ABE genes or large structural differences. We show a random set of allele-specific genes with a mean AD of 0.0249 as an example (**Figure 6B**, middle-right). MAE genes had a mean AD of 0.1748 (**Figure 6B**, right), significantly higher than randomly selected allelespecific genes (p < 0.01, Supplemental Methods E). This approach is useful for quantifying differences between maternal and paternal genomes throughout the cell cycle, highlighting gene sets with large structural or expression differences over time. In previous work, we have also shown that this method may be broadly applicable to time-series analysis of different cell types (*23*).

## Discussion

In this study, we present evidence for the intimate relationship among allele-specific gene expression, genome structure, and protein binding across the cell cycle. We validated our data and methods using known allele-specific properties such as the monoallelic expression of imprinted and X-linked genes, broad similarities of chromatin compartments between the maternal and paternal genomes, and large heterochromatic domains of Chromosome X (*9, 64–67*). Unique to this study, we established a coordination of allele-biased expression and changes in local genome structure, which included hundreds of genes not commonly associated with allele-biased expression. We observed further evidence of this coordination through corresponding protein binding biases.

Through our analysis of mature (nascent) RNA, we found 467 (380) genes to be differentially expressed between the two alleles and 229 (164) genes with differential expression through the cell cycle. Approximately half of the genes with allele-biased expression are only differentially expressed in certain cell cycle phases, and over half of the genes with CBE are only differentially expressed in one allele. Further research is needed to explore why certain genes have coordinated cell cycle dynamics across both alleles, while other genes have disparate expression in some cell cycle phases. We predict that these transient allelic biases may be associated with developmental pathologies and tumorigenesis, similar to imprinted and other MAE genes. Conversely, we found no allele-biased expression from genes in multiple biological modules, such as the cell cycle and glycolysis pathways (Table 1). We were not able to establish a statistical significance here due to the limited number of allele-specific genes in these modules, so we surveyed a set of 662 essential genes and found that they are significantly less likely to have allele-biased expression (*46*). This supports our hypothesis of highly coordinated biallelic expression in universally essential genes.

We developed a novel phasing algorithm, HaploHiC, which uses Hi-C reads mapped to phased SNVs/InDels to predict nearby reads of unknown parental origin. This allowed us to decrease the sparsity of our allele-specific contact matrices and increase confidence in our analysis of the parental differences in genome structure. While we found that the overall compartmentalization (euchromatin and heterochromatin) of the two genomes was broadly similar, there were many differences in TAD boundaries and local genome structure between the two genomes and between cell cycle phases. We focused our search for allele-specific differences in genome structure by calculating the similarity of local contacts surrounding differentially expressed genes (*23, 51*). We found that differentially expressed genes were significantly more likely to have corresponding changes in local genome structure than random allele-specific genes.

We incorporated publicly available allele-specific protein binding data for Pol II and CTCF to explore the mechanisms behind the gene expression and local genome structure relationship (*7*). In genes that had both allele-biased expression and Pol II binding biases, we found that 85% of these genes had allelic bias agreement. Additionally, we found that all of the genes with expression and Pol II binding bias agreement had significant changes in local genome structure. Analysis of the relationships among allele-specific gene expression, genome structure, and protein binding is currently hindered by the amount of information available and our limited understanding of the dynamics of cell-specific genome structure and gene expression variability (*68*). The ability to separate maternal and paternal gene expression and protein binding is dependent on the presence of a SNV/InDel within the gene body and nearby protein binding motifs. As SNVs/InDels are relatively rare in the human genome, the number of genes available to study is severely limited. Once we are able to separate the maternal and paternal genomes through advances in experimental techniques, we will be able to fully study these relationships.

Overall, these data support an intimate allele-specific relationship between genome structure and function, coupled through allele-specific protein binding. Changes in genome structure, influenced by the binding of proteins such as CTCF, can affect the ability of transcription factors and transcription machinery to access DNA. This results in changes in the rate of transcription of RNA, captured by Bru-seq. The rate of transcription leads to differential steady state gene expression, captured by RNA-seq. Integration of these data into a comprehensive computational framework led to the development of a maternal and paternal 4DN, which can be visualized using 4DN phase planes and quantified using allelic divergence. Allele-specific analysis across the cell cycle will be imperative to discern the underlying mechanisms behind many diseases by uncovering potential associations between deleterious mutations and allelic bias, and may have broad translational impact spanning cancer cell biology, complex disorders of growth and development, and precision medicine.

## Description of Supplemental Data

Supplemental Data include additional methods details, 11 figures, and 12 tables.

## Acknowledgements

We thank the University of Michigan Sequencing Core members for producing high quality Bru-seq, RNA-seq, and Hi-C data. We thank Michele Paulson and Dr. Mats Ljungman for generating the Bru-seq sequencing libraries and for helpful discussion. We also thank Dr. Jacob Kitzman for providing the GM12878 cell line and assisting in resolving genotype-phasing issues of the cell line. We thank Sam Dilworth and the iReprogram team for processing and formatting the KEGG pathway data. We are grateful to Dr. Thomas Ried for helpful reading of the manuscript. We thank the Air Force Office of Scientific Research (Award No: FA9550-18-1-0028), and the Forbes Institute for Cancer Discovery for supporting our work. G.S.O. acknowledges NIH grants P30ES0187885 and U24CA210967.

## Web Resources

Prototype MATLAB implementation for differential expression analysis can be found here: Identifying Differentially Expressed Genes from RNA-Seq Data. HaploHiC code is available through a GitHub repository:https://github.com/Nobel-Justin/HaploHiC.

## Data and Code Availability

The RNA-seq, Bru-seq, and Hi-C data used for this study are available through the Gene Expression Omnibus (GEO) database, https://www.ncbi.nlm.nih.gov/geo/ (accession number GSE159813).

## References

1. Alfred G Knudson. Mutation and cancer: statistical study of retinoblastoma. Proceedings of the National Academy of Sciences, 68(4):820–823, 1971.

2. Julius Gudmundsson, Gudrun Johannesdottir, Jon Thor Bergthorsson, Adalgeir Arason, Sigurdur Ingvarsson, Valgardur Egilsson, and Rosa Björk Barkardottir. Different tumor types from BRCA2 carriers show wild-type chromosome deletions on 13q12-q13. Cancer Research, 55(21):4830–4832, 1995.

3. Kara N Maxwell, Bradley Wubbenhorst, Brandon M Wenz, Daniel De Sloover, John Pluta, Lyndsey Emery, Amanda Barrett, Adam A Kraya, Ioannis N Anastopoulos, Shun Yu, et al. Brca locus-specific loss of heterozygosity in germline BRCA1 and BRCA2 carriers. Nature Communications, 8(1):1–11, 2017.

4. Irina S Zakharova, Alexander I Shevchenko, and Suren M Zakian. Monoallelic gene expression in mammals. Chromosoma, 118(3):279–290, 2009.

5. Karin Buiting. Prader–Willi syndrome and Angelman syndrome. In American Journal of Medical Genetics Part C: Seminars in Medical Genetics, volume 154, pages 365–376. Wiley Online Library, 2010.

6. 1000 Genomes Project Consortium. A global reference for human genetic variation. Nature, 526(7571):68, 2015.

7. Joel Rozowsky, Alexej Abyzov, Jing Wang, Pedro Alves, Debasish Raha, Arif Harmanci, Jing Leng, Robert Bjornson, Yong Kong, Naoki Kitabayashi, et al. AlleleSeq: analysis of allele-specific expression and binding in a network framework. Molecular Systems Biology, 7(1):522, 2011.

8. Anshul Kundaje, Wouter Meuleman, Jason Ernst, Misha Bilenky, Angela Yen, Alireza Heravi-Moussavi, Pouya Kheradpour, Zhizhuo Zhang, Jianrong Wang, Michael J Ziller, et al. Integrative analysis of 111 reference human epigenomes. Nature, 518(7539):317, 2015.

9. Suhas SP Rao, Miriam H Huntley, Neva C Durand, Elena K Stamenova, Ivan D Bochkov, James T Robinson, Adrian L Sanborn, Ido Machol, Arina D Omer, Eric S Lander, et al. A 3D map of the human genome at kilobase resolution reveals principles of chromatin looping. Cell, 159(7):1665–1680, 2014.

10. Longzhi Tan, Dong Xing, Chi-Han Chang, Heng Li, and X Sunney Xie. Three-dimensional genome structures of single diploid human cells. Science, 361(6405):924–928, 2018.

11. Danny Leung, Inkyung Jung, Nisha Rajagopal, Anthony Schmitt, Siddarth Selvaraj, Ah Young Lee, Chia-An Yen, Shin Lin, Yiing Lin, Yunjiang Qiu, et al. Integrative analysis of haplotype-resolved epigenomes across human tissues. Nature, 518(7539):350, 2015.

12. Indika Rajapakse and Mark Groudine. On emerging nuclear order. Journal of Cell Biology, 192(5):711–721, 2011.

13. Tom Misteli. The inner life of the genome. Scientific American, 304(2):66, 2011.

14. Peter R Cook. A model for all genomes: the role of transcription factories. Journal of Molecular Biology, 395(1):1–10, 2010.

15. Tom Misteli. The self-organizing genome: Principles of genome architecture and function. Cell, 2020.

16. Brian J Beliveau, Alistair N Boettiger, Maier S Avendaño, Ralf Jungmann, Ruth B Mc-Cole, Eric F Joyce, Caroline Kim-Kiselak, Frédéric Bantignies, Chamith Y Fonseka, Jelena Erceg, et al. Single-molecule super-resolution imaging of chromosomes and in situ haplotype visualization using oligopaint FISH probes. Nature Communications, 6(1):1–13, 2015.

17. Thomas Cremer and Marion Cremer. Chromosome territories. Cold Spring Harbor Perspectives in Biology, 2(3):a003889, 2010.

18. Joseph K Pickrell, John C Marioni, Athma A Pai, Jacob F Degner, Barbara E Engelhardt, Everlyne Nkadori, Jean-Baptiste Veyrieras, Matthew Stephens, Yoav Gilad, and Jonathan K Pritchard. Understanding mechanisms underlying human gene expression variation with RNA sequencing. Nature, 464(7289):768–772, 2010.

19. Jieming Chen, Joel Rozowsky, Timur R Galeev, Arif Harmanci, Robert Kitchen, Jason Bedford, Alexej Abyzov, Yong Kong, Lynne Regan, and Mark Gerstein. A uniform survey of allele-specific binding and expression over 1000-Genomes-Project individuals. Nature Communications, 7:11101, 2016.

20. Haiming Chen, Jie Chen, Lindsey A Muir, Scott Ronquist, Walter Meixner, Mats Ljungman, Thomas Ried, Stephen Smale, and Indika Rajapakse. Functional organization of the human 4D Nucleome. Proceedings of the National Academy of Sciences, 112(26):8002–8007, 2015.

21. Thomas Ried and Indika Rajapakse. The 4D nucleome. Methods (San Diego, Calif.), 123:1, 2017.

22. Job Dekker, Andrew S Belmont, Mitchell Guttman, Victor O Leshyk, John T Lis, Stavros Lomvardas, Leonid A Mirny, Clodagh C O’shea, Peter J Park, Bing Ren, et al. The 4D nucleome project. Nature, 549(7671):219–226, 2017.

23. Stephen Lindsly, Can Chen, Sijia Liu, Scott Ronquist, Samuel Dilworth, Michael Perlman, and Indika Rajapakse. 4DNvestigator: time series genomic data analysis toolbox. Nucleus, 12:1(58-64), 2021.

24. Erez Lieberman-Aiden, Nynke L Van Berkum, Louise Williams, Maxim Imakaev, Tobias Ragoczy, Agnes Telling, Ido Amit, Bryan R Lajoie, Peter J Sabo, Michael O Dorschner, et al. Comprehensive mapping of long-range interactions reveals folding principles of the human genome. Science, 326(5950):289–293, 2009.

25. Laura Seaman, Haiming Chen, Markus Brown, Darawalee Wangsa, Geoff Patterson, Jordi Camps, Gilbert S Omenn, Thomas Ried, and Indika Rajapakse. Nucleome analysis reveals structure-function relationships for colon cancer. Molecular Cancer Research, 15(7):821–830, 2017.

26. Michelle T Paulsen, Artur Veloso, Jayendra Prasad, Karan Bedi, Emily A Ljungman, Brian Magnuson, Thomas E Wilson, and Mats Ljungman. Use of Bru-Seq and BruChase-Seq for genome-wide assessment of the synthesis and stability of RNA. Methods, 67(1):45–54, 2014.

27. Thomas D Wu and Serban Nacu. Fast and SNP-tolerant detection of complex variants and splicing in short reads. Bioinformatics, 26(7):873–881, 2010.

28. Kimberly R Kukurba and Stephen B Montgomery. RNA sequencing and analysis. Cold Spring Harbor Protocols, 2015(11):pdb–top084970, 2015.

29. Jacob F Degner, John C Marioni, Athma A Pai, Joseph K Pickrell, Everlyne Nkadori, Yoav Gilad, and Jonathan K Pritchard. Effect of read-mapping biases on detecting allele-specific expression from RNA-sequencing data. Bioinformatics, 25(24):3207–3212, 2009.

30. Heng Li, Bob Handsaker, Alec Wysoker, Tim Fennell, Jue Ruan, Nils Homer, Gabor Marth, Goncalo Abecasis, and Richard Durbin. The sequence alignment/map format and SAM-tools. Bioinformatics, 25(16):2078–2079, 2009.

31. Simon Anders and Wolfgang Huber. Differential expression analysis for sequence count data. Genome Biology, 11(10):R106, 2010.

32. Yoav Benjamini and Yosef Hochberg. Controlling the false discovery rate: a practical and powerful approach to multiple testing. Journal of the Royal Statistical Society: Series B (Methodological), 57(1):289–300, 1995.

33. Siddarth Selvaraj, Jesse R Dixon, Vikas Bansal, and Bing Ren. Whole-genome haplotype reconstruction using proximity-ligation and shotgun sequencing. Nature Biotechnology, 31(12):1111, 2013.

34. Neva C Durand, Muhammad S Shamim, Ido Machol, Suhas SP Rao, Miriam H Huntley, Eric S Lander, and Erez Lieberman Aiden. Juicer provides a one-click system for analyzing loop-resolution Hi-C experiments. Cell Systems, 3(1):95–98, 2016.

35. Philip A Knight and Daniel Ruiz. A fast algorithm for matrix balancing. IMA Journal of Numerical Analysis, 33(3):1029–1047, 2013.

36. Irina Solovei, Antonio Cavallo, Lothar Schermelleh, Françoise Jaunin, Catia Scasselati, Dusan Cmarko, Christoph Cremer, Stanislav Fakan, and Thomas Cremer. Spatial preservation of nuclear chromatin architecture during three-dimensional fluorescence in situ hybridization (3D-FISH). Experimental Cell Research, 276(1):10–23, 2002.

37. Andreas Bolzer, Gregor Kreth, Irina Solovei, Daniela Koehler, Kaan Saracoglu, Christine Fauth, Stefan Müller, Roland Eils, Christoph Cremer, Michael R Speicher, et al. Threedimensional maps of all chromosomes in human male fibroblast nuclei and prometaphase rosettes. PLoS Biol, 3(5):e157, 2005.

38. Scott Ronquist, Walter Meixner, Indika Rajapakse, and John Snyder. Insight into dynamic genome imaging: Canonical framework identification and high-throughput analysis. Methods, 123:119–127, 2017.

39. Alexander Gimelbrant, John N Hutchinson, Benjamin R Thompson, and Andrew Chess. Widespread monoallelic expression on human autosomes. Science, 318(5853):1136–1140, 2007.

40. Qiaolin Deng, Daniel Ramsköld, Björn Reinius, and Rickard Sandberg. Single-cell RNA-seq reveals dynamic, random monoallelic gene expression in mammalian cells. Science, 343(6167):193–196, 2014.

41. Nuala A O’Leary, Mathew W Wright, J Rodney Brister, Stacy Ciufo, Diana Haddad, Rich McVeigh, Bhanu Rajput, Barbara Robbertse, Brian Smith-White, Danso Ako-Adjei, et al. Reference sequence (RefSeq) database at NCBI: current status, taxonomic expansion, and functional annotation. Nucleic Acids Research, 44(D1):D733–D745, 2016.

42. Yang Cao, Susan S AlHumaidi, Eissa A Faqeih, Beth A Pitel, Patrick Lundquist, and Umut Aypar. A novel deletion of SNURF/SNRPN exon 1 in a patient with Prader-Willi-like phenotype. European Journal of Medical Genetics, 60(8):416–420, 2017.

43. J Adams. Imprinting and genetic disease: Angelman, Prader-Willi and Beckwith–Weidemann syndromes. Nat Educ, 1(1), 2008.

44. Minoru Kanehisa and Susumu Goto. KEGG: kyoto encyclopedia of genes and genomes. Nucleic Acids Research, 28(1):27–30, 2000.

45. PT Wade Cohen, GS Omenn, AG Motulsky, S-H Chen, and ER Giblett. Restricted variation in the glycolytic enzymes of human brain and erythrocytes. Nature New Biology, 241(112):229–233, 1973.

46. Vincent A Blomen, Peter Májek, Lucas T Jae, Johannes W Bigenzahn, Joppe Nieuwenhuis, Jacqueline Staring, Roberto Sacco, Ferdy R van Diemen, Nadine Olk, Alexey Stukalov, et al. Gene essentiality and synthetic lethality in haploid human cells. Science, 350(6264):1092–1096, 2015.

47. Jesse R Dixon, Inkyung Jung, Siddarth Selvaraj, Yin Shen, Jessica E Antosiewicz-Bourget, Ah Young Lee, Zhen Ye, Audrey Kim, Nisha Rajagopal, Wei Xie, et al. Chromatin architecture reorganization during stem cell differentiation. Nature, 518(7539):331, 2015.

48. Jie Chen, Alfred O Hero III, and Indika Rajapakse. Spectral identification of topological domains. Bioinformatics, 32(14):2151–2158, 2016.

49. Jesse R Dixon, Siddarth Selvaraj, Feng Yue, Audrey Kim, Yan Li, Yin Shen, Ming Hu, Jun S Liu, and Bing Ren. Topological domains in mammalian genomes identified by analysis of chromatin interactions. Nature, 485(7398):376, 2012.

50. Elizabeth H Finn, Gianluca Pegoraro, Hugo B Brandao, Anne-Laure Valton, Marlies E Oomen, Job Dekker, Leonid Mirny, and Tom Misteli. Extensive heterogeneity and intrinsic variation in spatial genome organization. Cell, 176(6):1502–1515, 2019.

51. James A Koziol, Joel E Alexander, Lance O Bauer, Samuel Kuperman, Sandra Morzorati, Sean J O’connor, John Rohrbaugh, Bernice Porjesz, Henri Begleiter, and John Polich. A graphical technique for displaying correlation matrices. The American Statistician, 51(4):301–304, 1997.

52. Gordana Wutz, Csilla Várnai, Kota Nagasaka, David A Cisneros, Roman R Stocsits, Wen Tang, Stefan Schoenfelder, Gregor Jessberger, Matthias Muhar, M Julius Hossain, et al. Topologically associating domains and chromatin loops depend on cohesin and are regulated by CTCF, WAPL, and PDS5 proteins. The EMBO Journal, 36(24):3573–3599, 2017.

53. Nelle Varoquaux, Ferhat Ay, William Stafford Noble, and Jean-Philippe Vert. A statistical approach for inferring the 3D structure of the genome. Bioinformatics, 30(12):i26–i33, 2014.

54. Eyal Ben-David, Shahar Shohat, and Sagiv Shifman. Allelic expression analysis in the brain suggests a role for heterogeneous insults affecting epigenetic processes in autism spectrum disorders. Human Molecular Genetics, 23(15):4111–4124, 2014.

55. Fang-Ju Wu, Ting-Yu Lin, Li-Ying Sung, Wei-Fang Chang, Po-Chih Wu, and Ching-Wei Luo. BMP8A sustains spermatogenesis by activating both SMAD1/5/8 and SMAD2/3 in spermatogonia. Science Signaling, 10(477), 2017.

56. Yeawon Kim, Sun-Ji Park, Scott R Manson, Carlos AF Molina, Kendrah Kidd, Heather Thiessen-Philbrook, Rebecca J Perry, Helen Liapis, Stanislav Kmoch, Chirag R Parikh, et al. Elevated urinary CRELD2 is associated with endoplasmic reticulum stress–mediated kidney disease. JCI Insight, 2(23), 2017.

57. Joseph Petroziello, Andrew Yamane, Lori Westendorf, Melissa Thompson, Charlotte Mc-Donagh, Charles Cerveny, Che-Leung Law, Alan Wahl, and Paul Carter. Suppression subtractive hybridization and expression profiling identifies a unique set of genes overexpressed in non-small-cell lung cancer. Oncogene, 23(46):7734–7745, 2004.

58. Cameron S Osborne, Lyubomira Chakalova, Karen E Brown, David Carter, Alice Horton, Emmanuel Debrand, Beatriz Goyenechea, Jennifer A Mitchell, Susana Lopes, Wolf Reik, et al. Active genes dynamically colocalize to shared sites of ongoing transcription. Nature Genetics, 36(10):1065–1071, 2004.

59. Mark Newman. Networks. Oxford University Press, 2018.

60. Steven H Strogatz. Exploring complex networks. Nature, 410(6825):268–276, 2001.

61. Sijia Liu, Haiming Chen, Scott Ronquist, Laura Seaman, Nicholas Ceglia, Walter Meixner, Pin-Yu Chen, Gerald Higgins, Pierre Baldi, Steve Smale, et al. Genome architecture mediates transcriptional control of human myogenic reprogramming. iScience, 6:232–246, 2018.

62. Josée Dostie, Todd A Richmond, Ramy A Arnaout, Rebecca R Selzer, William L Lee, Tracey A Honan, Eric D Rubio, Anton Krumm, Justin Lamb, Chad Nusbaum, et al. Chromosome conformation capture carbon copy (5c): a massively parallel solution for mapping interactions between genomic elements. Genome Research, 16(10):1299–1309, 2006.

63. Lawrence Page, Sergey Brin, Rajeev Motwani, and Terry Winograd. The PageRank citation ranking: Bringing order to the web. Technical report, Stanford InfoLab, 1999.

64. Wolf Reik and Jörn Walter. Genomic imprinting: parental influence on the genome. Nature Reviews Genetics, 2(1):21, 2001.

65. Tomas Babak, Brian DeVeale, Emily K Tsang, Yiqi Zhou, Xin Li, Kevin S Smith, Kim R Kukurba, Rui Zhang, Jin Billy Li, Derek van der Kooy, et al. Genetic conflict reflected in tissue-specific maps of genomic imprinting in human and mouse. Nature Genetics, 47(5):544, 2015.

66. Yael Baran, Meena Subramaniam, Anne Biton, Taru Tukiainen, Emily K Tsang, Manuel A Rivas, Matti Pirinen, Maria Gutierrez-Arcelus, Kevin S Smith, Kim R Kukurba, et al. The landscape of genomic imprinting across diverse adult human tissues. Genome Research, 25(7):927–936, 2015.

67. Federico A Santoni, Georgios Stamoulis, Marco Garieri, Emilie Falconnet, Pascale Ribaux, Christelle Borel, and Stylianos E Antonarakis. Detection of imprinted genes by single-cell allele-specific gene expression. The American Journal of Human Genetics, 100(3):444–453, 2017.

68. Elizabeth H Finn and Tom Misteli. Molecular basis and biological function of variability in spatial genome organization. Science, 365(6457), 2019.

