## Supplemental Materials for "Functional Organization of the Maternal and Paternal Human 4D Nucleome"

### Supplemental Methods

#### A Generation and Haplotype Assignment of Hi-C

##### A.1 Phased germline mutations of GM12878

From the AlleleSeq database (version: Jan-7-2017) (*1*), we downloaded the VCF file of sample GM12878. This VCF file contains phased heterozygous germline mutations (SNV and InDel) with a total of 2.49 million genomic loci (**Table S8**). We confirmed that paternal allele is the left side of phased alleles, and the maternal allele is the right side, e.g., '0|1' at HG19 genomic position chr1:2276371. This mutation list was also utilized in the realignment process of GATK.

HaploHiC utilizes heterozygous genotype information to distinguish reads' parental origin as long as the reads cover the heterozygous genomic loci. Considering that short sequence insertions and deletions dramatically influence alignment accuracy, we applied a filtration on heterozygous InDels. All homologous and heterozygous InDel alleles were assigned a minimum distance to the next InDel on the haplotyped genome. This minimum distance considers repeatability of both the mutation sequence and local genomic context. The mutation sequence (i.e., the inserted or deleted sequence) might be repetitive and repeated at its genomic position. For example, at HG19 genomic position chr1:2277268, the maternal genome has a deletion ('CACA'). The deleted sequence is 'CA'-unit repetitive (2 times) and also repeated in the adjacent reference context ('CA'-unit repeats 11 times). The minimum distance is the whole repeats' length added by unit repeated time of mutation sequence. So, at genomic position chr1:2277268, the minimum distance of the deletion allele on maternal genome is 28, which means any InDel located within this distance on maternal genome will be excluded from following analysis. In total, 11,614 heterozygous genomic loci harboring InDel alleles were filtered (**Table S8**). HaploHiC outputs a list of these filtered genotypes. The minimum distance is also applied in InDel allele judgment on sequencing reads (see A.3).

##### A.2 Alignment and filtering of Hi-C PE-reads

Illumina adapter sequences and low quality ends were trimmed from raw Hi-C reads by Trimmomatic (*2*). Paired-ends (PE, R1 and R2) were separately aligned to the human reference genome (HG19) using the function ‘mem’ from BWA (0.7.15) with default parameters (*3*). Then, each alignment BAM file was coordinated, sorted, and realigned by GATK (3.5) (*4*) with the VCF file described in **Supplemental Methods A.1**. All realigned BAM files were sorted by read names before processing in HaploHiC.

HaploHiC loads Hi-C PE-reads’ mapping information from paired alignment BAM files, and filters unqualified and invalid pairs in the first step. After exclusion of unqualified and invalid pairs, remaining Hi-C PE-reads are eligible to provide chromatin contacts (**Table S9**).

Unqualified PE-reads are filtered out if either or both ends are un-mapped, mapped with a mapping quality less than 20, or mapped to multiple genomic positions. Note that supplementary alignments are reserved, because theoretically, a portion of sequencing reads that cover ligation site could be soft-clipped mapped to two genomic locations by BWA and provide contact information. HaploHiC sets two filtering criteria on primary and supplementary alignments: 1) the difference between their length sum and the reads’ length must be less than one fifth of the reads’ length, 2) the overlaps between primary and supplementary mapped parts must be shorter than one third of the shorter one. If the two criteria are not satisfied simultaneously, the supplementary alignment will be marked as ‘remove_SP’ and discarded.

HaploHiC detects four categories of invalid pairs: 1) dangling ends, 2) self circle, 3) dumped pair forwarded mapped, and 4) dumped pair reversed mapped. The definition of these four categories are the same as HiC-Pro (*5*), while the singleton type is included in the unqualified PE-reads mentioned above. The two ends of an invalid pair must map to same the enzyme fragment or within a close distance (1 kb).

##### A.3 Parental origin categories of Hi-C PE-reads

HaploHiC assigns a haplotype situation to sequencing reads by checking whether reads cover and support heterozygous alleles of specific parental origin. Note that, in this section, we deal with each alignment of sequencing reads. Four haplotype situations are defined (**Figure 4**): 1) 'P' (only paternal allele), 2) 'M' (only maternal allele), 3) 'I' (both maternal and paternal alleles, i.e., inter-haplotype), and 4) 'U' (no parental allele supported). Clearly, the former three situations ('P', 'M', and 'I') are haplotype-known, and the fourth ('U') is haplotype-unknown. Note that sequencing reads with 'I' support both haplotypes simultaneously, as covering the junction site of the ligated maternal and paternal fragments.

For sequencing reads covering heterozygous genomic positions and matching either parental allele, the bases that match the allele must meet two criteria: 1) base quality is not less than 20, and 2) distance to bilateral edges of this read is not less than 5 bp. Note, if the allele is determined through an InDel, the read-edge distance might use the minimum distance representing the repeatability mentioned above (see section A.1) if the latter is larger. HaploHiC outputs a list recording all InDel alleles that have the repeatability distance. There are a total of 255,961 heterozygous loci having such InDel alleles in sample GM12878 (**Table S8**).

According to the haplotype status of the two ends, HaploHiC assigns Hi-C PE-reads to seven categories: dEnd-P/M/I, sEnd-P/M/I, and dEnd-U. Here, 'dEnd' and 'sEnd' represent dual-ends and single-end, respectively. Common instances of each category are shown in **Figure 4**. Additionally, we also set requirements on distance of alignments (**Table S9**).

- dEnd-P: both ends support only paternal alleles, and are not close-aligned
- dEnd-M: both ends support only maternal alleles, and are not close-aligned
- dEnd-I: at least one end supports both maternal and paternal alleles, and the other one is haplotype-known
- sEnd-P: one end supports only paternal allele, the other one has haplotype-unknown alignment

- sEnd-M: one end supports only maternal allele, the other one has haplotype-unknown alignment
- sEnd-I: one end supports both maternal and paternal alleles, the other one has haplotype-unknown alignment
- dEnd-U: both ends are haplotype-unknown.

Clearly, four categories (dEnd-P/M/I and sEnd-I) have confirmed allele-specific contacts, and we call them phased Hi-C PE-reads. Conversely, the other three categories are unphased: PE-reads have one (sEnd-P/M) or two (dEnd-U) haplotype-unknown ends. The next step is to assign the haplotype to these unphased Hi-C PE-reads based on the allele-specific contact information in local regions.

##### A.4 Local region contacts from phased Hi-C PE-reads

By integrating all phased Hi-C PE-reads, HaploHiC records allele-specific contacts in windowed regions. For example, to record allele-specific contacts of two genomic regions ('no.A' window on 'chrA', and 'no.B' window on 'chrB'), HaploHiC sorts 'chrA' and 'chrB' in ASCII order, and if 'chrA' and 'chrB' are the same chromosome, it then sorts 'no.A' and 'no.B' in ascending order. This dual-sorting process avoids duplicated records and reduces software memory consumption. After sorting, the former region is 'chrF, no.F' (here, 'F' for former), and the latter region is 'chrL, no.L' (here, 'L' for latter). HaploHiC keeps a dictionary with key-value pairs. The key is 'chrF, no.F, chrL, no.L', and its value is a set of phased Hi-C PE-reads that link these two regions with haplotype combinations (i.e., 'P-P', 'M-M', 'P-M', and 'M-P', **Figure 4B**, **Figure S4**). Additionally, HaploHiC de-duplicates the Hi-C PE-reads under each haplotype combination.

HaploHiC utilizes a local allele-specific contacts based algorithm to impute the haplotype for haplotype-unknown ends. To get the local region of one end of the unphased Hi-C PE-reads, HaploHiC extends bilaterally from the mapped position. The extension length is 50 kb on each unilateral side, which is referred to as an extension unit. Note that each unilateral region cannot harbor more than 30 phased heterozygous loci, or else the unilateral region will be trimmed. The bilateral extended regions are merged as one local region. A similar extension gets the local region of the other end of the Hi-C PE-read. Then, from the dictionary of phased Hi-C pairs, HaploHiC counts PE-reads linking these two local regions of each haplotype combination. For simplicity, the contact counts of haplotype combinations are: $\alpha$ ('P-P'), $\beta$ ('M-M'), $\gamma$ ('P-M'), and $\delta$ ('M-P'), respectively. HaploHiC uses these counts to impute the haplotype for haplotype-unknown reads. Note that if the sum of these counts is zero, local regions of both ends will be iteratively extended by more extension units until the sum is non-zero or local regions reach the maximum length (1 Mb). Finally, if the sum is still zero, we define this pair of local regions as unphased.

##### A.5 Assign haplotype to unphased Hi-C PE-reads (sEnd-P/M)

First, HaploHiC deals with unphased Hi-C PE-reads from sEnd-P/M categories. Because these Hi-C pairs already have one haplotype-known end, some haplotype combinations should be excluded. For example, one Hi-C pair has one paternal end (mapped to 'posA' on 'chrA') and one haplotype-unknown end (mapped to 'posB' on 'chrB'). After dual-sorting the mapped chromosomes and positions, 'chrB, posB' is the 'chrF, posF', and 'chrA, posA' is the 'chrL, posL'. HaploHiC calculates the local regions of 'chrF, posF' and 'chrL, posL' respectively, and summarizes contacts counts of haplotype combinations recorded under key 'chrF, posF, chrL, no.L': $\alpha$ ('P-P'), $\beta$ ('M-M'), $\gamma$ ('P-M'), and $\delta$ ('M-P'). Because 'chrL, posL' is from the paternal genome in this instance, $\beta$ ('M-M') and $\gamma$ ('P-M') are impossible and should be excluded. HaploHiC then randomly assigns 'P' or 'M' to the haplotype-unknown end ('chrF, posF') with possibility defined as:

$$possibility(paternal) = \frac{\alpha}{\alpha + \delta} \tag{1}$$

$$possibility(maternal) = \frac{\delta}{\alpha + \delta} \tag{2}$$

Moreover, if the local regions are still unphased after iterative extension, i.e., the sum of contacts counts of haplotype combinations is still zero, HaploHiC will assign the haplotype with uniform possibility depending on the mapped chromosome situation. In this example, if 'chrF, posF' and 'chrL, posL' belong to same chromosome, HaploHiC assigns the haplotype of 'chrL, posL' (the haplotype-known end) to 'chrF, posF' (the haplotype-unknown end). However, if 'chrF, posF' and 'chrL, posL' belong to different chromosomes, uniform possibility (0.5) will be applied.

If intra-chromosome mapped (in this example):

$$possibility(paternal) = 1 \tag{3}$$

$$possibility(maternal) = 0 \tag{4}$$

If inter-chromosome mapped:

$$possibility(paternal) = 0.5 \tag{5}$$

$$possibility(maternal) = 0.5 \tag{6}$$

Hi-C pairs from sEnd-P/M categories are marked as 'phased imputed' and 'unphased imputed' corresponding to phased and unphased local regions, respectively.

##### A.6 Assign haplotype to unphased Hi-C PE-reads (dEnd-U)

Before the operations on unphased Hi-C PE-reads from dEnd-U category, HaploHiC records 'phased imputed' Hi-C pairs from sEnd-P/M categories to expand the contacts dictionary. For dEnd-U Hi-C pairs, calculation of local contacts counts of haplotype combinations is identical to that of sEnd-P/M mentioned above. Note that as both ends are haplotype-unknown, no haplotype combination will be excluded. Based on local regions' contacts count ($\alpha$ ('P-P'), $\beta$ ('M-M'), $\gamma$ ('P-M'), and $\delta$ ('M-P'), **Figure 4B**), HaploHiC randomly assigns a haplotype combination to dEnd-U Hi-C PE-reads with possibility defined as:

$$possibility(paternal, paternal) = \frac{\alpha}{\alpha + \beta + \gamma + \delta} \tag{7}$$

$$possibility(maternal, maternal) = \frac{\beta}{\alpha + \beta + \gamma + \delta} \tag{8}$$

$$possibility(paternal, maternal) = \frac{\gamma}{\alpha + \beta + \gamma + \delta} \tag{9}$$

$$possibility(maternal, paternal) = \frac{\delta}{\alpha + \beta + \gamma + \delta} \tag{10}$$

Similar to sEnd-P/M, if the local regions are still unphased after iterative extension, HaploHiC will assign a haplotype with uniform possibility depending on the mapped chromosome status.

If intra-chromosome mapped:

$$possibility(paternal, paternal) = 0.5 \tag{11}$$

$$possibility(maternal, maternal) = 0.5 \tag{12}$$

$$possibility(paternal, maternal) = 0 \tag{13}$$

$$possibility(maternal, paternal) = 0 \tag{14}$$

If inter-chromosome mapped:

$$possibility(paternal, paternal) = 0.25 \tag{15}$$

$$possibility(maternal, maternal) = 0.25 \tag{16}$$

$$possibility(paternal, maternal) = 0.25 \tag{17}$$

$$possibility(maternal, paternal) = 0.25 \tag{18}$$

Hi-C pairs from dEnd-U category are also marked as 'phased imputed' and 'unphased imputed' corresponding to phased and unphased local regions, respectively.

##### A.7 Allele-specific integrated results of Hi-C PE-reads

After processing the unphased Hi-C PE-reads of sEnd-P/M and dEnd-U categories, HaploHiC successfully assigns haplotype to all valid Hi-C pairs. Finally, HaploHiC integrates Hi-C pairs of each haplotype combination: intra-paternal ('P-P'), intra-maternal ('M-M'), and inter-haplotype ('P-M' and 'M-P'). Note that 'P-M' and 'M-P' are recorded with different tags in one file.

- Intra-paternal includes dEnd-P category, and imputed ('P-P') Hi-C pairs from sEnd-P/M and dEnd-U categories
- Intra-maternal includes dEnd-M category, and imputed ('M-M') Hi-C pairs from sEnd-P/M and dEnd-U categories

- Inter-haplotype includes dEnd-I and sEnd-I categories, and imputed ('P-M' and 'M-P') Hi-C pairs from sEnd-P/M and dEnd-U categories

All integrated files are in BAM format, which keeps the original alignments of Hi-C pairs. HaploHiC adds several tags in SAM optional fields to denote processing details. A report is generated recording statistics of each category of all Hi-C pairs.

We calculated the phased rate in each sample (**Table S10**). The phased rate is the percentage of phased Hi-C pairs, including dEnd-P/M/I, sEnd-I, and phased Hi-C pairs with imputed haplotype from sEnd-P/M and dEnd-U categories. Phased rates in sEnd-P/M and dEnd-U categories are calculated separately.

Additionally, we introduced 'inter-chr imbalance' to evaluate the difference between inter-haplotype and intra-haplotype assignment of inter-chromosome Hi-C pairs. Theoretically, for inter-chromosome contacts, there is no reason to assume any difference between the intra-haplotype and inter-haplotype. The 'inter-chr imbalance' is defined as the difference of intra-haplotype and inter-haplotype inter-chromosome contacts divided by their larger one. Our data shows very low 'inter-chr imbalance' (0.01%-0.04%, **Table S10**), which supports the accuracy of HaploHiC phasing.

##### A.8 Validation of allele-specific contacts

To validate HaploHiC, we randomly removed 10% of heterozygous loci from the list of phased mutations. The Hi-C PE reads from three categories (dEnd-P/M/I) were selected for validation, as both ends of these reads have known parental origin and can be used as the ground truth. To estimate the imputation accuracy of HaploHiC, we compared the imputed haplotype and the original haplotype of all Hi-C PE that cover the removed phased mutations. This validation method is similar to the one proposed in Tan *et. al.* (*6*). We found that HaploHiC was able to correctly assign an average of 96.9%, 97.2%, and 97.3% of these Hi-C reads for G1, S, and G2, respectively, over 10 trials. The minimum accuracy over all trials for G1, S, and G2 were 96.8%, 97.1%, and 97.3%. Accuracy of imputation was calculated by the fraction of correctly imputed ones from the haplotype-known Hi-C reads covering these removed heterozygous mutations (**Table S12**).

To further evaluate our local contacts based algorithm, we simulated Hi-C sequencing data from haplotype specific contacts between gene pairs of ten categories:
Five categories for imitation of intra-haplotype autosome contacts:

- Cross-Chrom: inter-chromosome translocations
- Long-Distance: intra-chromosome, but on different arms
- Long-Distance: intra-chromosome, on same arm, gene distance is $>$ 10 Mb
- TAD level: intra-chromosome, on same arm, gene distance is [1 Mb, 2 Mb]
- LOOP level: intra-chromosome, on same arm, gene distance is $<$ 700 kb

Two categories for imitation of inter-haplotype contacts:

- InterHap/interChr: inter-chromosome translocations
- InterHap/intraChr: intra-chromosome, gene distance is $>$ 10 Mb

Three categories for imitation of chrX specific activation (intra-haplotype):

- chrX/LongDistance: gene distance is $>$ 10 Mb
- chrX/TAD: gene distance is [1 Mb, 2 Mb]
- chrX/LOOP: gene distance falls into is $<$ 700 kb

In total, 67 gene pairs were randomly selected from COSMIC cancer gene census database (*7*) to construct pairwise allele-specific contacts of maternal and paternal genomes respectively (**Table S12**). The maternal and paternal genomes are downloaded from AlleleSeq database (version: Jan-7-2017) (*1*). The simulations on the LOOP level are CN-based (copy number), and the others are SV-based (structure variation, **Figure S5**). To imitate SV-based chromatin contacts for one pair of genes, e.g. *BCR* and *ABL1*, breakpoints are randomly picked from the two genes' genomic regions respectively. At the breakpoints, reciprocal translocations are formed via concatenating extended flanking 2Mb genomic sequences. As the gene-pair contact is heterozygous, sequences are extracted from paternal genome FASTA file to form an allele-specific SV, and sequences from maternal genome are extracted and kept unchanged (**Figure S5A**). To simulate CN-based chromatin contacts on the LOOP level, we assign different copy numbers to genomic regions harboring two neighbor genes on the two haplotypes (**Figure S5B**). For one genomic region containing two nearby genes (gene *C* and *D*), to make the paternal genome have more contacts between these two genes, we use two copies of this region from the paternal genome, while only keeping one copy from the maternal genome. We applied simu3C (*8*) to simulate the Hi-C sequencing reads of the SV-based and CN-based allele-specific gene-pair contacts.

HaploHiC successfully recovers the allele-specific contacts in simulation data with 97.66% accuracy (**Table S12**). First, all SV-based intra-haplotype allele-specific contacts are precisely reported by HaploHiC. For example, contacts of gene pair '*ETV6*, *NTRK3*' are all 'P-P', and contacts of gene pair '*KCNJ5*, *LMO2*' are all 'M-M'. Second, for CN-based intra-haplotype allele-specific contacts, the advantage haplotype is successfully reported by HaploHiC. For example, gene pair '*FLT3*, *LNX2*' has 'P-P' contacts more than three times of the 'M-M' contacts, and gene pair '*RUNX1*, *SMIM11*' has 'M-M' contacts more than two times of the 'P-P' contacts. These ratios are close to what we found in simulation. Third, all inter-haplotype gene pairs are successfully identified with correct the haplotype combination. For example, all contacts of the gene pair '*WT1*, *ZMYM2*' link paternal *WT1* and maternal *ZMYM2*. The gene pair '*MAP2K1*, *NUP214*' has seven 'M-P' contacts (false positives), which is only 2.6% of all its contacts (273). Fourth, all the intra-haplotype chrX allele-specific activation cases are reported correctly. Two gene pairs have some bias to inactivated haplotype ('*LAS1L*, *ZC4H2*' and '*GPC3*, *HS6ST2*'). Note that even shifted, they still show enrichment on correct haplotype. Gene pair '*LAS1L*, *ZC4H2*' has larger shifting (33.3%) to 'M-M' contact, because the neighbor gene (*MSN*, gene distance is about 75 kb) of *LAS1L* forms gene pair '*MSN*, *STAG2*' which is simulated to have only 'M-M' contacts. '*LAS1L*, *ZC4H2*' is influenced by '*MSN*, *STAG2*' as we merged all simulated Hi-C sequencing data of 67 gene pairs together as one sample in HaploHiC evaluation. Finally, all Hi-C pairs with mistakenly assigned haplotype combinations are gathered as false positives (count is 1,326, total count of valid Hi-C pairs is 55,013).

#### B Whole Chromosome Probe Generation and 3D FISH

Whole chromosome paint probes were generated in-house using PCR labeling techniques as described at https://ccr.cancer.gov/Genetics-Branch/thomas-ried. Chromosome 7 was labeled with Orange dUTP (Abbott Laboratories, Abbott Park, IL), Chromosome 8 was labeled with Dy505 (Dyomics, Jena, Germany) and Chromosome 11 was labeled with Biotin-16-dUTP (Roche Applied Science, Indianapolis, IL). Cells were grown on slides and fixed with 4% paraformaldehyde for 10 minutes. Cells were then washed with 0.05% Triton X100 for five minutes followed by permeabilization steps which included incubation with 0.5% Triton X100 for 20 minutes, followed by subsequent repeated (4x) freeze thaw in liquid nitro- gen/glycerol. The slides were then incubated in 20% glycerol for at least one hour before being frozen in 1xPBS at -20°C until hybridization was performed. Prior to hybridization, cells were washed in 0.05% Triton X100 followed by incubation in 0.1N HCl for 10 minutes. Cells were then washed in 2XSSC followed by incubation in 50% formamide/2XSSC for at least one hour before hybridization. Cells and probes were co-denatured at 72°C for five minutes followed by a 48 hours hybridization at 37°C. After incubation at 37°C, detection commences with posthybridization washes followed by incubation in blocking (3% BSA, 4xSSC, 0.1% Tween20) for 30 minutes at 37°C. The biotinylated probes were detected with the fluorochrome Cy5 conjugated to Streptavidin (Rockland, Gilbertsville, PA). The slides were then washed with 2XSSC before being counterstained with Prolong Gold antifade reagent with DAPI (Promega Madison, WI).

#### C Spectral clustering: The Laplacian, Fiedler value, and Fiedler vector

The Laplacian, Fiedler value, and Fiedler vector can be summarized as follows. Consider an adjacency matrix $\mathbf{A}$, where $(\mathbf{A})_{i,j} = w(n_i, n_j)$, and weight function, $w$, satisfying $w(n_i, n_j) = w(n_j, n_i)$ (symmetrical) and $w(n_i, n_j) \geq 0$ (nonnegative). The Laplacian, $\mathbf{L}$, of $\mathbf{A}$ is defined to be $\mathbf{L} = \mathbf{D} - \mathbf{A}$, where $\mathbf{D} = diag(d_1, \cdots, d_k)$ and $d_i = \Sigma_{j=1}^{k} a_{ij}$. The normalized Laplacian is the matrix $\overline{\mathbf{L}} = \mathbf{D}^{-1/2}\mathbf{L}\mathbf{D}^{-1/2}$. The second smallest eigenvalue of $\mathbf{L}$ (or $\overline{\mathbf{L}}$) is called the Fiedler value, and the corresponding eigenvector is called the Fiedler vector (*9*). The Fiedler value is also known as the algebraic connectivity of a graph. The magnitude of the Fiedler value increases as the number of edges in the graph increases, and as the graph becomes more structurally ordered. The Fiedler vector partitions the genome into two parts that reflect underlying topology, as given by edge weights inferred from Hi-C data. The Fiedler vector plays a role similar to the eigenvector associated with the largest eigenvalue (principal component 1) of the correlation matrix of the normalized Hi-C matrix (*10*), but it is directly related to properties of the associated graph (*9*). The Fiedler vector can be calculated recursively to identify smaller partitions in Hi-C data. These smaller partitions correspond to topologically associated domains (TADs) (*11*). Description of the Laplacian, Fiedler value, and Fiedler vector adapted from (*12*).

#### D Structure-Function Visualization and the 4D Nucleome

The goal of the maternal and paternal 4DN is to understand the relationship of allele-specific genomic structure and gene function through time. Gene expression (RNA-seq) directly offers us a scalar value to represent the function of an individual gene, or for a genomic locus (total gene expression for all genes within the locus). In order to find a compatible scalar value which represents structure, we look to Hi-C contacts. Hi-C provides insight into the structural organization of the genome by finding distant regions of the genome (in terms of genomic sequence) that are close to one another in 3D space. The 3D structure of the genome can be viewed through the perspective of a ‘genomic network’. In such a network, genomic regions are considered nodes and the contacts between genomic regions are the edges.

A concept that is well known in network theory is centrality. Network centrality measurements, or features, encompass a wide range of network properties (*13*). The common goal of these features is to assign quantitative measurements to the structure of the network. One particularly important feature is the eigenvector centrality, which is the eigenvector associated with the largest eigenvalue of the adjacency matrix which defines the network. Eigenvector centrality assigns values to each node in a network corresponding to that node's influence on the network. Other centrality features that have been shown to have biological relevance include degree, betweenness, and closeness centrality (*14*). We calculated these four centrality features from the Hi-C data at 1 Mb resolution and concatenated them with RNA-seq to form a new structure-function (S-F) matrix which represents both structure and function (rows correspond to genomic loci, columns correspond to centrality features and RNA-seq) (*14, 15*). We derived the S-F matrix both genome-wide and for individual chromosomes. We then normalize the S-F matrix for each setting (maternal and paternal in G1, S, and G2) and concatenate all settings. Next, we apply t-SNE to the combined S-F matrix (containing all settings) and reduce it to two dimensions (*16*). Then we can visualize the two dimensional projection, and observe how the maternal and paternal genomes compare across cell cycle phases (**Figure S9**).

The structure-function matrix integrates genomic structure and gene expression into a common subspace, but it does not directly inform us of the relationship between structure and function. To visualize this allele-specific relationship, we use eigenvector centrality and mature RNA (RNA-seq, FPKM) for each allele at each phase of the cell cycle. In other words, eigenvector centrality and RNA-seq serve as structure ($x$-axis) and function ($y$-axis) coordinates across the cell cycle, respectively (**Figure 6**). The cell cycle and B cell receptor signaling genes displayed in **Figure 6** are contained in a larger sub-network of genes based on their KEGG pathway, whose Hi-C contacts (1 Mb resolution) define the edges of their respective sub-networks (from which eigenvector centrality is calculated). Similarly, the MAE genes displayed are contained within a sub-network of all MAE genes identified in our ABE analysis. The allele-specific genes displayed, which serve as a control, are contained within a gene sub-network which was randomly selected from all allele-specific genes. The number of allele-specific genes selected for the control is the mean of the other three sets' sizes. The three points (G1, S, and G2) in the 4DN phase plane for each allele are fit with a minimum volume ellipse to capture the 4DN variance of the allele (*17*). Allelic divergence for each gene is calculated as the mean Euclidean distance between the maternal and paternal alleles' coordinates in the 4DN phase plane for G1, S, and G2, after normalization of coordinates. The allelic divergence for each group of genes displayed in **Figure 6** is defined as the mean of these genes' individual allelic divergences.

#### E Statistical Significance via Permutation Test

A permutation test builds the shape of null hypothesis (namely, the random background distribution) by resampling the observed data. We use a permutation test to establish statistical significance in our analysis of the relationship between local structural changes and corresponding functional changes, the frequency of ABE genes in an essential gene set, and average allelic divergence of MAE genes. This sampling procedure was repeated 10,000 times in all cases cases. A rank-based p-value is then calculated for the right-tailed event for testing likelihood of local structure changes around ABE genes and allelic divergence. The left-tailed event was tested for decreased ABE in an essential gene set (*18*). The background distribution for ABE in essential genes was generated by calculating the proportion of ABE genes in a randomly selected set of 662 allele-specific genes (same number of genes as the allele-specific essential gene set). The proportion of ABE genes in the essential gene set compared to this background distribution yields the p-value (**Figure S3**). The background distributions for genome structure changes were generated by calculating the average number of significant changes in the Hi-C contacts surrounding random allele-specific genes. The probability of the right-tailed event of our observation of significant structural changes around ABE, CBE, and combined set of ABE and CBE genes under their respective background distributions yields the p-values (**Figure S7**). The background distribution for allelic divergence in MAE genes was generated by calculating the average allelic divergence among randomly selected sets of allele-specific genes. The allelic divergence of the MAE gene set compared to this background distribution yields the p-value (**Figure S11**).

### F Structure Alignment and 3D Modeling

Structures computed from different distance matrices could be varied in both scale and orientation. To align different structures and superimpose them into the same coordinates, we used Procrustes analysis (*19*) which applies the optimal transform to the second matrix (including scaling/dilation, rotations, and reflections) to minimize the sum of square errors of the point-wise differences. First, we translocate shapes to the origin by subtracting the mean value of all coordinates. Next, we force shapes into the same scale by dividing each shape by the Frobenius norm. For an $m \times n$ matrix $\mathbf{A}$, the Frobenius norm is defined as $\|\mathbf{A}\|_F = (\sum_{i=1}^{m} \sum_{j=1}^{n} |a_{ij}|^2)^{\frac{1}{2}}$. Finally, we find an optimal rotation matrix that will align one contact matrix $\mathbf{A}$ to matrix $\mathbf{B}$ by using Singular Value Decomposition (SVD) on $\mathbf{M}$ where $\mathbf{M} = \mathbf{A}^{\top}\mathbf{B}$. Applying SVD to the matrix $\mathbf{M}$ gives $\mathbf{M} = \mathbf{U}\mathbf{\Sigma}\mathbf{V}^{\top}$, where $\mathbf{V}^{\top}$ is the rotation matrix of $\mathbf{B}$. Then $\mathbf{B}\mathbf{V}^{\top}$ is the rotated shape. To visualize the final structure from the inferred three dimensional embedding, we smooth the curve by interpolating three dimensional scatter data using the radial basis function (RBF) kernel. The structure was visualized using the Mayavi package in Python (*20*).

### Supplemental Figures

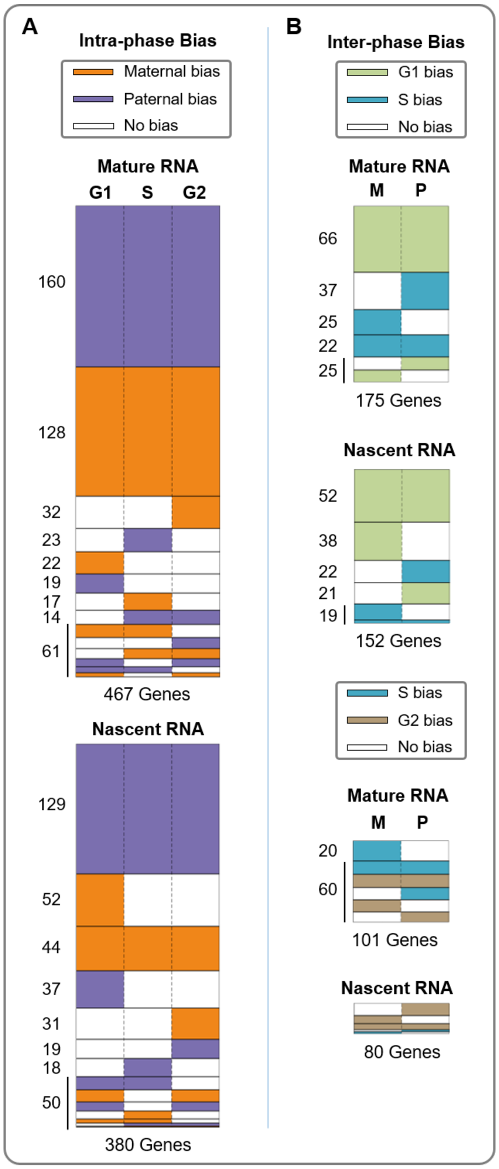


Fig. S1: Expression biases identified between alleles and cell cycle phases. **(A)** Differential expression between the maternal and paternal alleles for each cell cycle phase. Differentially expressed genes with a bias towards the maternal (paternal) allele in a particular cell cycle phase are shaded orange (purple). Each row represents a group of genes with a specific expression bias pattern. Analogous to **Figure 3**. **(B)** Differential expression between cell cycle phases within each allele. Top shows differential expression between G1 and S phases while bottom shows differential expression between S and G2. Genes are shaded in the color of the cell cycle phase in which their expression is significantly higher. G1, S, and G2 are shaded green, blue, brown respectively. In both (A) and (B), white represents no expression bias.

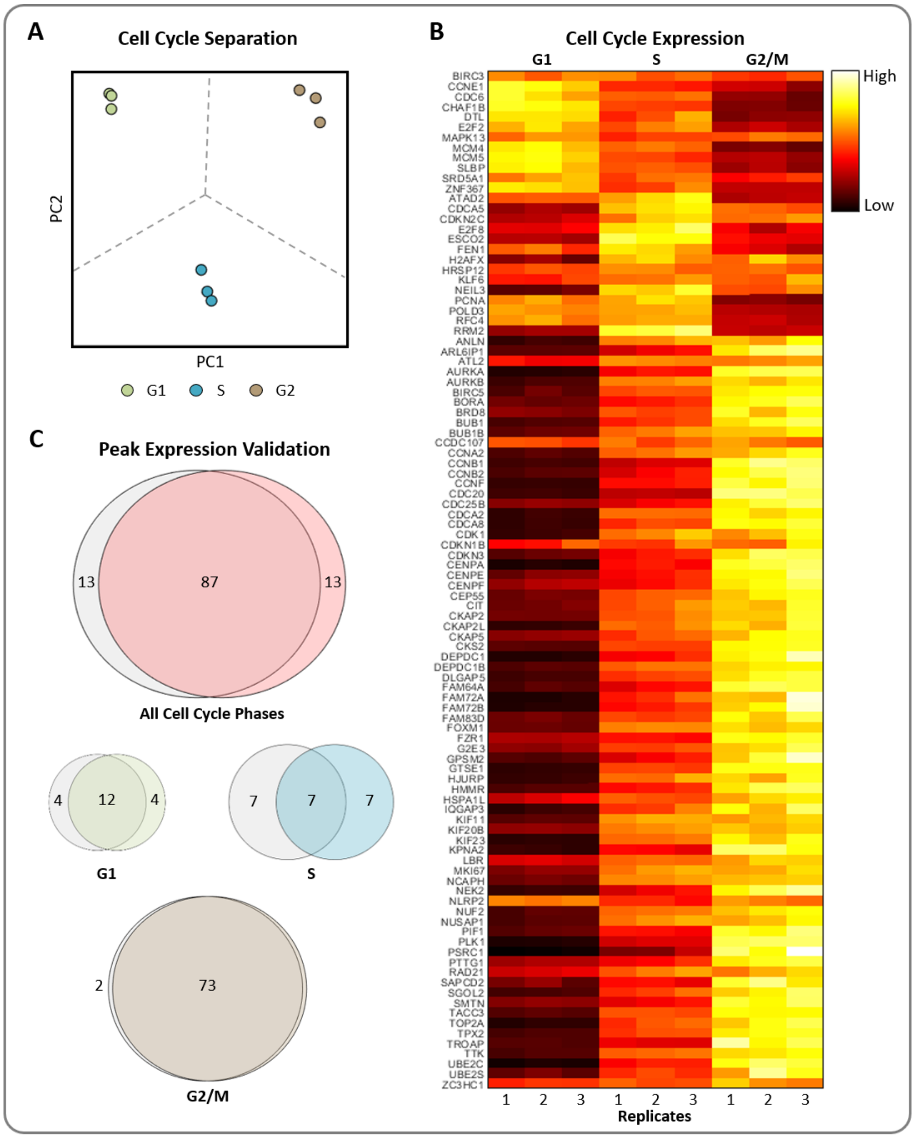


Fig. S2: Cell cycle separation of RNA-seq data. **(A)** Principal component analysis (PCA) of three replicates over three time points. G1, S, and G2 are easily separated and the replicates of each cell cycle phase cluster after dimension reduction. **(B)** RNA-seq expression heatmap of the top 100 periodically expressed genes in our data from Cyclebase are row normalized and clustered by peak expression time (*21*). **(C)** Venn diagram comparison for the peak expression time of the same 100 periodically expressed genes. Red, green, blue, and brown circles represent the peak expression times for G1, S, and G2 within our data respectively, and overlap with gray indicates agreement with Cyclebase peak expression times. Over all phases, 87% of these genes match in their peak expression cell cycle phase.

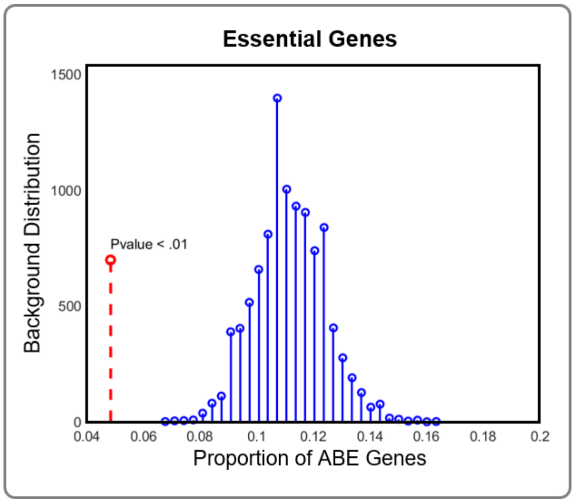


Fig. S3: Statistical significance of decreased ABE in essential genes. A permutation test was performed to determine whether essential genes were less likely to have ABE versus randomly sampled allele-specific genes (**Supplemental Methods E**).

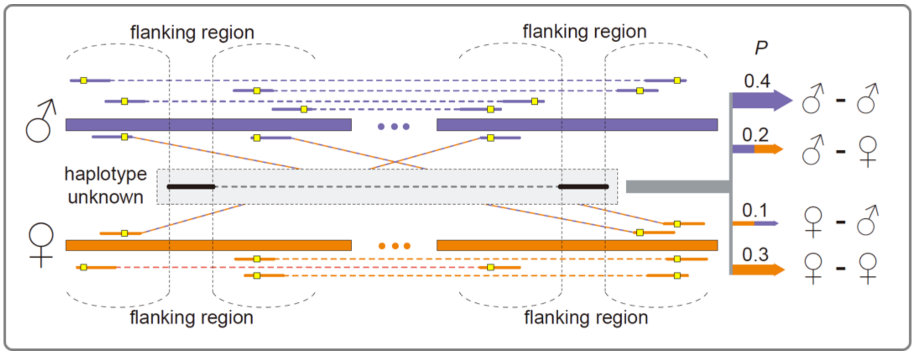


Fig. S4: Expanded diagram of HaploHiC phasing in **Figure 4**. Haplotype-unknown Hi-C pair is distributed to certain haplotype combination with probabilities depending on local phased contacts in flanking regions. Homologous genomic regions are linked by phased Hi-C pairs (purple for paternal, orange for maternal). Phased heterozygous loci are denoted by yellow squares.

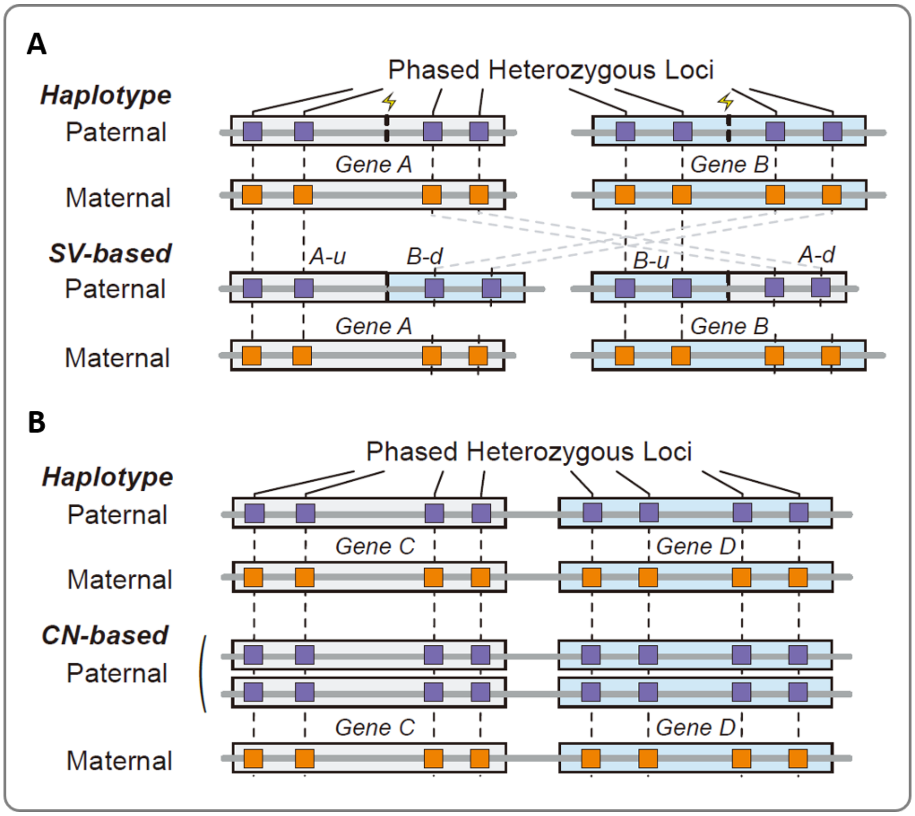


Fig. S5: Simulation of SV-based and CN-based contacts. **(A)** Gene pairs are reciprocally partially concatenated on the paternal haplotype to form a chimeric sequence. The maternal haplotype carries the wild-type. Upstream and downstream genomic regions are denoted by 'u' and 'd', respectively. **(B)** The copy number of gene pairs' genomic region on the paternal haplotype is increased, and the maternal haplotype is kept at one copy.

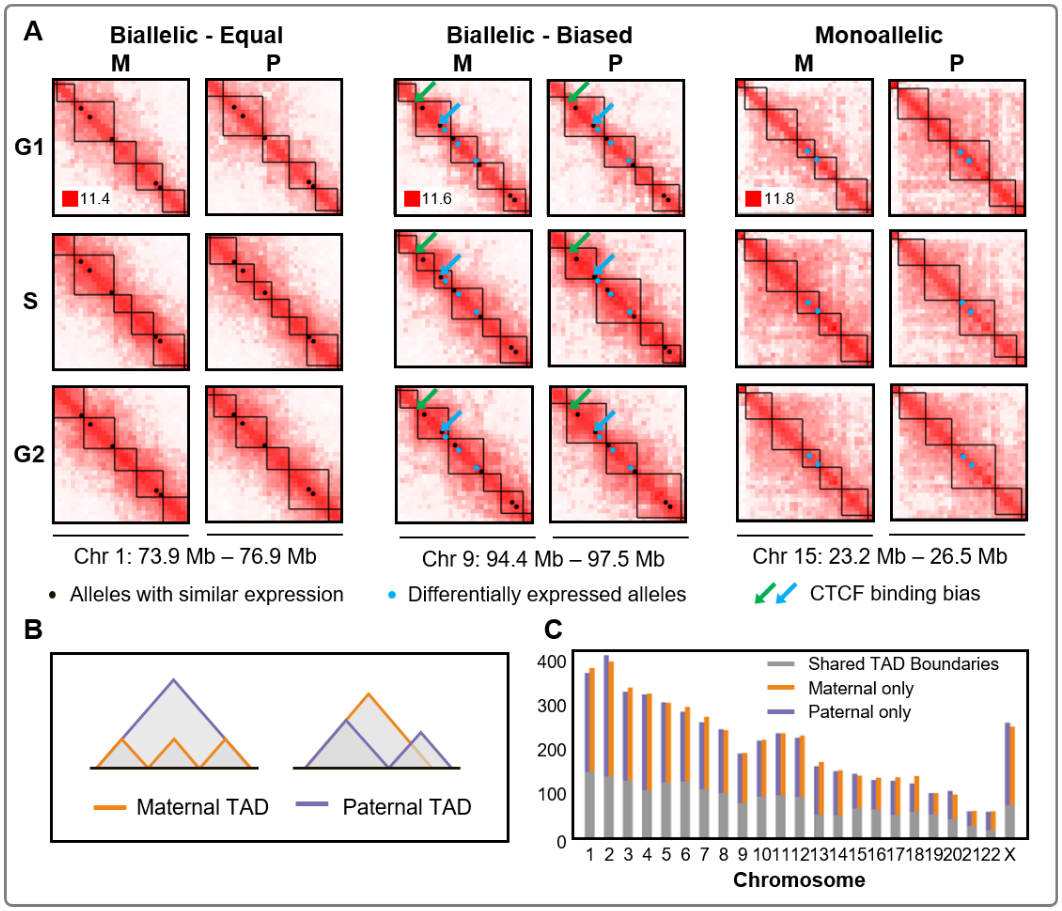


Fig. S6: Allele-specific topologically associated domains (TADs). (**A**) Subsection of Chromosomes 1 (left), 9 (middle), and 15 (right) separated into allele- and cell cycle-specific matrices shown in $log_2$ scale 100 kb resolution, with TADs depicted as solid black lines. Circles in each setting represent allele-specific genes with similar (black) and differential (blue) expression. Arrows indicate a binding bias of CTCF to the maternal (green) or paternal (blue) genome (*1*). The CTCF binding data is not cell cycle separated. Arrows were added to all cell cycle phases, but CTCF binding may only be active at particular phases. TAD boundaries are found using the spectral identification method (*11*). **(B)** Schematic representation of allele-specific TADs. A single TAD in one genome can be identified as a multiple TADs in the other genome (left), or TAD boundaries can shift depending on local genome structure (right). **(C)** Bar graph highlighting the number of TAD boundaries in common between maternal and paternal (gray), maternal only TAD boundaries (orange), and paternal only TAD boundaries (purple) for all chromosomes in G1.

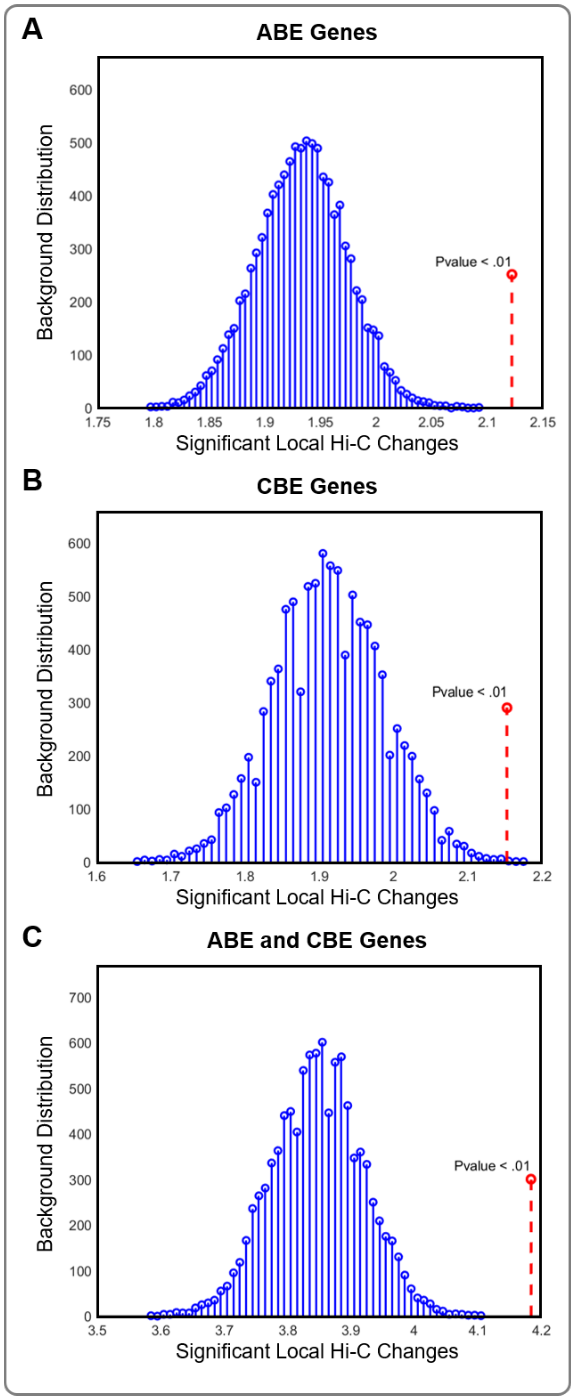


Fig. S7: Statistical significance of changes in chromatin structure around differentially expressed genes. Permutation tests were performed for ABE genes, CBE genes, and the combined set of ABE and CBE genes to determine whether significant local structure changes were more likely in differentially expressed genes versus randomly sampled allele-specific genes for their respective comparisons (**Supplemental Methods E**).

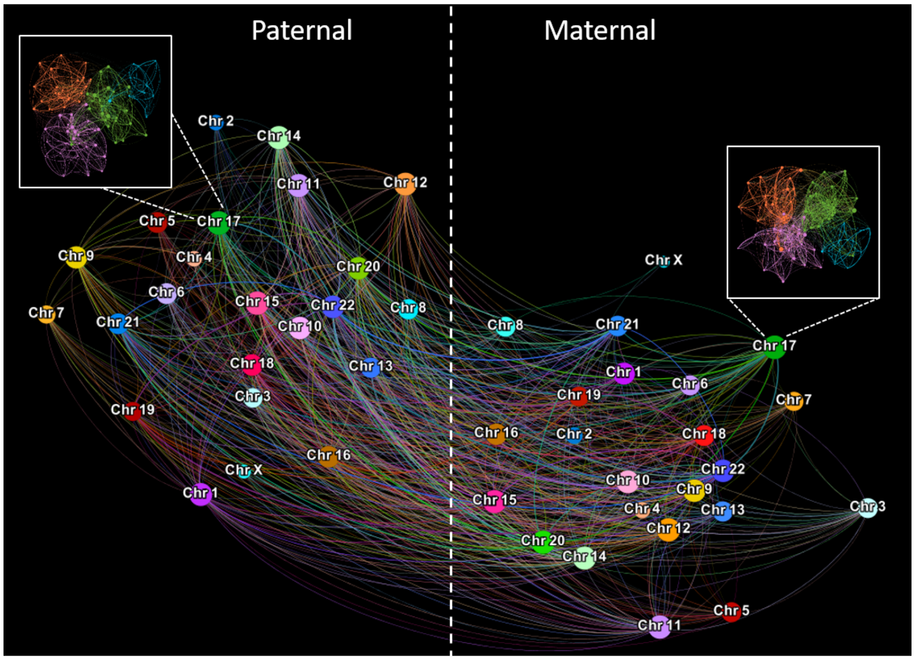


Fig. S8: Maternal and paternal genomes as a network of chromosomes. The maternal and paternal chromosomes are separated to highlight inter-haplotype interactions from Hi-C data. Each chromosome has an internal network of intra-chromosomal chromatin contacts as highlighted in the maternal and paternal Chromosome 17.

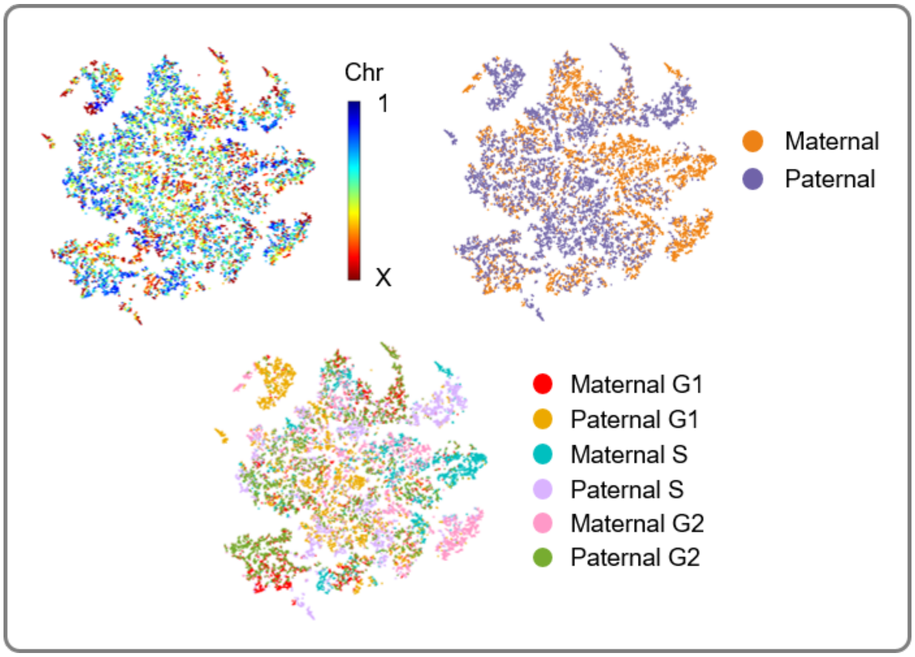


Fig. S9: Genome-wide visualization of the maternal and paternal 4DN. Low dimensional representation (t-SNE) of structure and function for the maternal and paternal genomes across cell cycle phases (1 Mb resolution) (**Supplemental Methods D**) (*16*). The three plots show the same data with different color schemes to highlight the locations of chromosomes (top left), parental origin (top right), and parental origin with cell cycle phase (bottom). Differences in the low dimensional mappings can be seen genome-wide for the maternal and paternal genomes, as well as cell cycle phases.

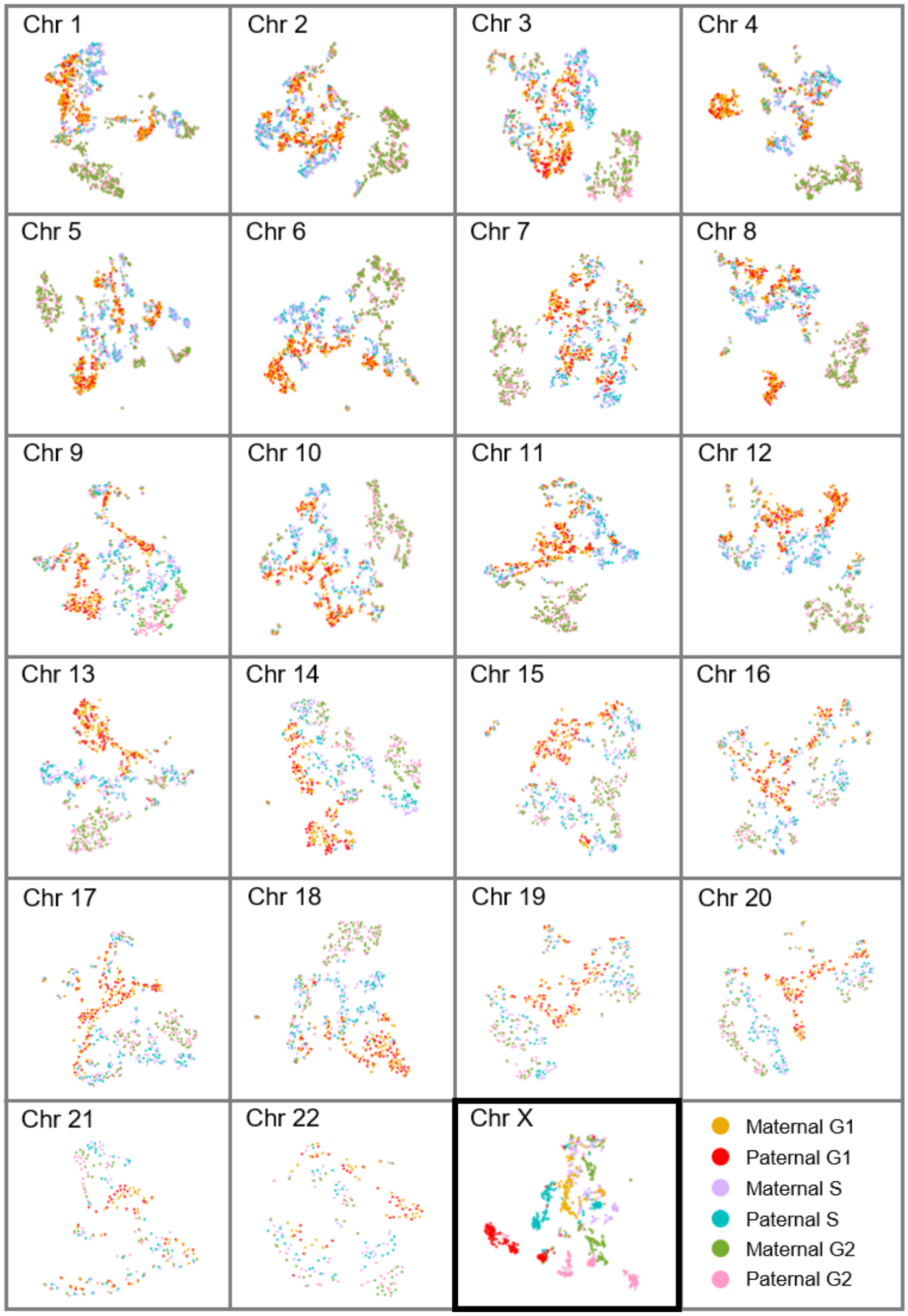


Fig. S10: The same process from **Figure S9** was applied to individual chromosomes, with color scheme highlighting parental origin and cell cycle phase. The X Chromosome shows clear separation of the maternal and paternal copies. The autosomes have more subtle differences between their maternal and paternal low dimensional projections.

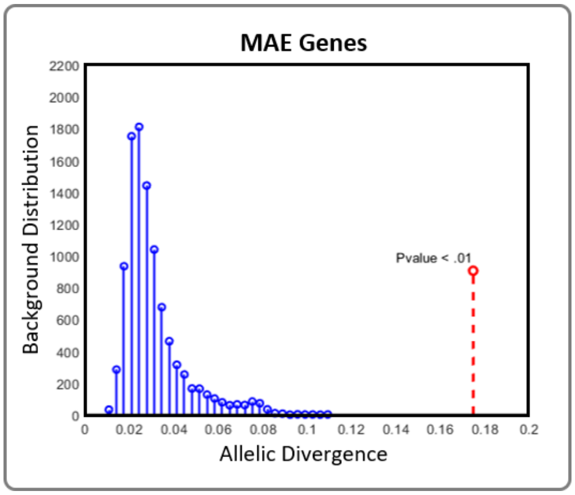


Fig. S11: Statistical significance of allelic divergence in monoallelicly expressed genes. A permutation test was performed for a subset of monoallelicly expressed genes to determine whether their allelic divergence was significantly higher than randomly sampled allele-specific genes (**Supplemental Methods E**).

#### Supplemental Tables

Table S1: Differential expression per chromosome. Number of allele-specific genes, differentially expressed genes between alleles (ABE) and cell cycle phases (CBE), and percentage of allele-specific genes that are differentially expressed for each chromosome.

Table S2: Differential expression between alleles using RNA-seq. ABE was observed for 467 genes between the maternal and paternal alleles in mature RNA.

Table S3: Differential expression between alleles using Bru-seq. ABE was observed for 380 genes between the maternal and paternal alleles in nascent RNA.

Table S4: Differential expression between cell cycle phases using RNA-seq. CBE was observed for 229 genes between cell cycle phases for the maternal or paternal alleles in mature RNA.

Table S5: Differential expression between cell cycle phases using Bru-seq. CBE was observed for 164 genes between cell cycle phases for the maternal or paternal alleles in nascent RNA.

Table S6: Coordination of allele-biased TF binding and gene expression. Genes are separated into three groups for Pol II, CTCF, and other TFs. Each gene contains the number of binding instances, the parental bias, and whether or not the bias had agreement with their ABE.

Table S7: Allele biased expression of KEGG pathways. All KEGG pathway gene sets with at least five allele-specific genes were examined for ABE. Here, we list the the percentage of allele-specific genes with ABE for each pathway.

Table S8: Details of phased germline mutations of GM12878. Mutation counts of diverse allele scenarios on maternal and paternal genomes are listed.

Table S9: Parental origin categories of Hi-C PE-reads in HaploHiC. Seven Hi-C pairs categories are denoted with judgment conditions of paired-end alignments.

Table S10: Details of Hi-C PE-reads in sample GM12878 processed by HaploHiC. Hi-C pair counts of all categories in three steps are listed.

Table S11: Details of ten categories of gene pairs in Hi-C simulation and evaluation of HaploHiC. In total, 67 gene pair details are listed with parental and maternal origin contacts division results from HaploHiC.

Table S12: Details of SNV holdout validation of HaploHiC. Validation was performed independently on Hi-C data from G1, S, and G2 and repeated over 10 trials.

### References


1. Joel Rozowsky, Alexej Abyzov, Jing Wang, Pedro Alves, Debasish Raha, Arif Harmanci, Jing Leng, Robert Bjornson, Yong Kong, Naoki Kitabayashi, et al. Alleleseq: analysis of allele-specific expression and binding in a network framework. *Molecular systems biology*, 7(1):522, 2011.

2. Anthony M Bolger, Marc Lohse, and Bjoern Usadel. Trimmomatic: a flexible trimmer for illumina sequence data. *Bioinformatics*, 30(15):2114–2120, 2014.

3. Heng Li and Richard Durbin. Fast and accurate short read alignment with burrows–wheeler transform. *bioinformatics*, 25(14):1754–1760, 2009.

4. Aaron McKenna, Matthew Hanna, Eric Banks, Andrey Sivachenko, Kristian Cibulskis, Andrew Kernytsky, Kiran Garimella, David Altshuler, Stacey Gabriel, Mark Daly, et al. The genome analysis toolkit: a mapreduce framework for analyzing next-generation dna sequencing data. *Genome research*, 20(9):1297–1303, 2010.

5. Nicolas Servant, Nelle Varoquaux, Bryan R Lajoie, Eric Viara, Chong-Jian Chen, Jean-Philippe Vert, Edith Heard, Job Dekker, and Emmanuel Barillot. Hic-pro: an optimized and flexible pipeline for hi-c data processing. *Genome biology*, 16(1):259, 2015.

6. Longzhi Tan, Dong Xing, Chi-Han Chang, Heng Li, and X Sunney Xie. Three-dimensional genome structures of single diploid human cells. *Science*, 361(6405):924–928, 2018.

7. Simon A Forbes, David Beare, Harry Boutselakis, Sally Bamford, Nidhi Bindal, John Tate, Charlotte G Cole, Sari Ward, Elisabeth Dawson, Laura Ponting, et al. Cosmic: somatic cancer genetics at high-resolution. *Nucleic acids research*, 45(D1):D777–D783, 2016.

8. Matthew Z DeMaere and Aaron E Darling. Sim3c: simulation of hi-c and meta3c proximity ligation sequencing technologies. *GigaScience*, 7(2):gix103, 2017.

9. Fan RK Chung and Fan Chung Graham. *Spectral graph theory*. Number 92 in CBMS Regional Conference Series in Mathematics. American Mathematical Soc., 1997.

10. Erez Lieberman-Aiden, Nynke L Van Berkum, Louise Williams, Maxim Imakaev, Tobias Ragoczy, Agnes Telling, Ido Amit, Bryan R Lajoie, Peter J Sabo, Michael O Dorschner, et al. Comprehensive mapping of long-range interactions reveals folding principles of the human genome. *science*, 326(5950):289–293, 2009.

11. Jie Chen, Alfred O Hero III, and Indika Rajapakse. Spectral identification of topological domains. *Bioinformatics*, 32(14):2151–2158, 2016.

12. Haiming Chen, Jie Chen, Lindsey A Muir, Scott Ronquist, Walter Meixner, Mats Ljungman, Thomas Ried, Stephen Smale, and Indika Rajapakse. Functional organization of the human 4d nucleome. *Proceedings of the National Academy of Sciences*, 112(26):8002–8007, 2015.

13. Mark Newman. *Networks*. Oxford university press, 2018.

14. Sijia Liu, Haiming Chen, Scott Ronquist, Laura Seaman, Nicholas Ceglia, Walter Meixner, Pin-Yu Chen, Gerald Higgins, Pierre Baldi, Steve Smale, et al. Genome architecture mediates transcriptional control of human myogenic reprogramming. *iScience*, 6:232–246, 2018.

15. Stephen Lindsly, Can Chen, Sijia Liu, Scott Ronquist, Samuel Dilworth, Michael Perlman, and Indika Rajapakse. 4DNvestigator: time series genomic data analysis toolbox. *Nucleus*, 12:1(58-64), 2021.

16. Laurens van der Maaten and Geoffrey Hinton. Visualizing data using t-sne. *Journal of machine learning research*, 9(Nov):2579–2605, 2008.

17. P. Sun and R. M. Freund. Computation of minimum-volume covering ellipsoids. *Operations Research*, 52(5):690–706, 2004.

18. Vincent A Blomen, Peter Májek, Lucas T Jae, Johannes W Bigenzahn, Joppe Nieuwenhuis, Jacqueline Staring, Roberto Sacco, Ferdy R van Diemen, Nadine Olk, Alexey Stukalov, et al. Gene essentiality and synthetic lethality in haploid human cells. *Science*, 350(6264):1092–1096, 2015.

19. Mikkel B Stegmann and David Delgado Gomez. A brief introduction to statistical shape analysis. *Informatics and mathematical modelling, Technical University of Denmark, DTU*, 15(11), 2002.

20. Prabhu Ramachandran and Gaël Varoquaux. Mayavi: 3d visualization of scientific data. *Computing in Science & Engineering*, 13(2):40–51, 2011.

21. Alberto Santos, Rasmus Wernersson, and Lars Juhl Jensen. Cyclebase 3.0: a multi-organism database on cell-cycle regulation and phenotypes. *Nucleic acids research*, 43(D1):D1140–D1144, 2015.
